# Intronic polyadenylation isoforms in the 5’ part of genes constitute a source of microproteins and are involved in cell response to cisplatin

**DOI:** 10.1101/2023.12.05.569446

**Authors:** Alexandre Devaux, Iris Tanaka, Mandy Cadix, Amélie Heneman-Masurel, Sophie Michallet, Quentin Fouilleul, Alina Chakraborty, Céline M. Labbe, Nicolas Fontrodona, Jean-Baptiste Claude, Marc Deloger, Pierre Gestraud, Ludovic Tessier, Hussein Mortada, Sonia Lameiras, Virginie Raynal, Sylvain Baulande, Nicolas Servant, Didier Auboeuf, Béatrice Eymin, Stéphan Vagner, Martin Dutertre

## Abstract

Transcript isoforms generated by intronic polyadenylation (IPA) are widely regulated in various biological processes and often encode protein isoforms. Microproteins are small proteins translated from small open reading frames (sORFs) in noncoding RNAs and mRNAs, but their production by IPA isoforms is unknown. Using 3’-seq and long-read RNA-seq analyses in lung cancer cells, we show that cisplatin, a DNA-crosslinking anticancer agent, upregulates IPA isoforms relative to full-length mRNAs in long genes. A subset of cisplatin-regulated IPA isoforms are poorly associated with heavy polysomes and terminate upstream of the annotated translation initiation codon of genes. Such IPA isoforms in the *PHF20* and *PRKAR1B* genes are associated with light polysomes, contain Ribo-Seq-supported sORFs in an alternative last exon within the annotated 5’UTR part of genes, and are translated into microproteins. For *PRKAR1B*, the microprotein was detected by Western blot and immunofluorescence after transfection of a tagged isoform; and siRNA depletion of the endogenous IPA isoform, CRISPR deletion of the IPA site, or CRISPR mutation of the sORF initiation codon led to increased cell survival to cisplatin. Based on Ribo-Seq and mass-spectrometry data sets, we identified 156 genes producing both a canonical protein-coding mRNA and a microprotein-coding 5’UTR-located IPA isoform (coined miP-5’UTR-IPA isoform) regulated by cisplatin. Finally, the regulation of (miP-5’UTR-)IPA *versus* full-length isoforms by cisplatin involved an inhibition of transcription processivity in a FANCD2 and senataxin-dependent manner. Altogether, these findings reveal the novel paradigm of miP-5’UTR-IPA genes and their role in cancer cell response to a genotoxic agent.

**HIGHLIGHTS:**

- Cisplatin increases intronic-polyadenylation *versus* full-length transcript isoforms in long genes through a FANCD2 and senataxin-dependent decrease of transcription processivity
- A subset of cisplatin-regulated intronic-polyadenylation isoforms terminate in the annotated 5’UTR part of genes and encode microproteins, thus we coined them miP-5’UTR-IPA isoforms
- The miP-5’UTR-IPA isoform of *PRKAR1B* impacts cisplatin sensitivity and its effect is mediated by its small ORF
- We identify 156 genes producing both a canonical protein-coding mRNA and a microprotein-coding miP-5’UTR-IPA transcript

## INTRODUCTION

Intronic polyadenylation (IPA), sometimes referred to as splicing-dependent, upstream region- or coding region-alternative polyadenylation (APA), corresponds to the use of a polyadenylation site (polyA site) located upstream of the last exon (LE) of a gene and allows the production of transcripts with alternative last exons (ALEs) (1–4). Genome-wide analyses using 3’-seq (RNA-seq focused on the 3’ end of polyadenylated transcripts) and other approaches have revealed regulation of the IPA:LE isoform ratio in many human genes across tissues and in various biological processes, such as cell differentiation, proliferation, and cell responses to DNA-damaging agents and other stress inducers, with different stressors inducing different global patterns of IPA:LE isoform regulation (5–13). The mechanisms of IPA regulation have been intensively studied, with many regulatory factors acting at the levels of splicing, polyadenylation, and transcription elongation and processivity (1–4).

In contrast, relatively little is known about the translation outcome of IPA isoforms. On the one hand, dozens of IPA isoforms were shown to encode protein isoforms (3) and a recent genome-wide analysis showed that the majority of cytosolic IPA isoforms are associated with polysomes, suggesting they are translated (11). On the other hand, some studies identified two IPA isoforms with noncoding RNA functions (8, 14), IPA isoforms degraded by the nuclear exosome (15, 16), and abundant IPA isoforms with a low predicted coding potential that were therefore classified as long noncoding RNAs (lncRNAs), but their function and translation status were not studied (17). Microproteins, also called micropeptides or sORF-encoded peptides, are an emerging class of proteins of less than 100 amino acids that are encoded by small open reading frames (sORFs) found in lncRNAs, canonical mRNAs (either upstream, downstream, overlapping or inside the annotated canonical ORF), and circular RNAs (18, 19). An increasing number of human microproteins are involved in biological processes (18, 20, 21). IPA isoforms constitute a potential but unexplored source of microprotein production.

Cisplatin is a DNA-crosslinking anticancer agent that is widely used in non-small cell lung cancer (NSCLC) treatment (22). It widely regulates alternative splicing (23, 24) but little is known about its effects on IPA isoforms. In this study, by investigating on IPA isoform regulation by cisplatin in NSCLC cells, we identified cisplatin-regulated IPA isoforms that terminate within the annotated 5’UTR part of genes, encode microproteins, and impact cell response to cisplatin. These findings reveal the existence of genes (which we call miP-5’UTR-IPA genes) producing both a canonical protein-coding mRNA and a microprotein-coding IPA transcript.

## MATERIAL AND METHODS

### Plasmid construction

*PHF20* sORF#1, *PRKAR1B* sORF#1 and *PRKAR1B* sORF#2, together with the ALE sequence upstream of the sORF and with a carboxy-terminal Flag tag, were obtained by RT-PCR (using primers described in Table S1) and cloned into the pTRIPΔU3-MND-IRES-GFP plasmid (25) between the MND promoter and an internal ribosome entry site driving the expression of green fluorescent protein (GFP).

### Cell culture, transfection, and treatment

Cells were cultured in RPMI-1640-Glutamax (H358) or DMEM (A549, HeLa, HEK-293T) medium (GibcoBRL, Life Technologies, Cergy Pontoise, France) supplemented with 10% (v/v) heat-inactivated fetal calf serum (GibcoBRL), in 5% CO_2_ at 37°C. All cell lines and CRISPR clones were authenticated by STR analysis. Cisplatin was obtained from Selleckchem (Euromedex, Souffelweyersheim, France) and resuspended in DMSO. Unless otherwise stated, cisplatin was used at 25 and 100 µM in A549 and H358 cells, respectively. Cell treatment with vehicle (DMSO) was always done in parallel to cisplatin treatment. Reverse transfection of siRNAs (Eurogentec, Belgium; Table S2) was performed using Lipofectamine RNAimax (ThermoFischer Scientific, France) and following the manufacturer’s instructions. For plasmid transfection, cells were seeded in 6-well plates containing Marienfeld Superior cover glasses; the next day, cells were transfected for 24 hours with 5 µg of plasmid, 7.5 µL of Lipofectamine 3000 and 10 µL of P3000 (ThermoFischer Scientific, France) in the case of HeLa cells, or with 4 µg of plasmid and 9 µL of Lipofectamine 2000 (ThermoFischer Scientific, France) in the case of HEK-293T cells.

### WST1 and FACS analysis

For cell growth and viability analysis, cells were seeded in 96-well plates. Following cisplatin treatment, cell viability was assayed in triplicate wells using WST1 (Sigma Aldrich, France) according to the manufacturer’s instructions. For FACS analysis of DNA content, cells were fixed with 70% cold ethanol for 30 min on ice, treated with RNase A (20 µg/ml) for 20 min and stained with propidium iodide (10 µg/ml). Flow cytometric analysis of 10000 cells was performed on a FACScan flow cytometer (BD Biosciences) and data were recovered using the CellQuest software (BD Biosciences).

### Protein extraction and Western blot

For immunoblotting, cells washed three times in PBS were lysed in RIPA buffer (150mM NaCl, 50mM Tris HCl pH 8, 0.1% SDS, 1% Nonidet P40, 0.5% Na deoxycholate, 0.1mM PMSF, 2.5μg/ml pepstatin, 10μg/ml aprotinin, 5μg/ml leupeptin, 0.2mM Na_3_VO_4_) for 30 min on ice and pelleted. Protein concentration was determined using the Pierce BCA Protein Assay kit (Biorad). Proteins were then separated in 10-12% SDS-PAGE gels (or Tris-Tricine gel 16% gels for microproteins) and electroblotted onto PVDF membranes. Membranes were incubated overnight at +4°C with antibodies against PRIM2 (PA5-48859; ThermoFisher Scientific), PRKAR1B (PA5-55392, ThermoFischer Scientific), actin (Sigma-Aldrich, Lyon, France), GAPDH (G8795, Merck) or Flag (F1804, Sigma Aldrich), then for one hour with a horse-radish peroxidase-conjugated goat anti-mouse or anti-rabbit antibody (Jackson ImmunoResearch Laboratories, West Grove, PA, USA). After washing, blots were revealed using the ECL chemiluminescence method (Amersham, Les Ulis, France), according to the manufacturer’s protocol.

### Immunofluorescence analysis

Transfected cells grown on Marienfeld Superior cover glasses were washed twice with ice-cold PBS, fixed using 4% paraformaldehyde in PBS during 20 min, washed three times with PBS, permeabilized for 10 min with PBS-0.1% Triton-X, washed twice with PBS, blocked for 5 min with PBS-5% BSA, incubated with Anti-Flag M2 antibody (F1804, Sigma Aldrich) at RT for 1 hour, washed twice with PBS, blocked again, incubated with F(ab’)2-rabbit anti-mouse IgG (H+L) cross-adsorbed secondary antibody, Alexa Fluor 594 (A-21205, Invitrogen) in PBS-5% BSA for 1 hour, washed twice with PBS, incubated at RT with PBS containing 4’,6-diamidino-2-phenylindole (DAPI) 0.1 µg/mL, and washed twice with PBS. Cover glasses were mounted on slides in PBS, glycerol 15%, 1.4-diazabicyclo-(2.2.2) octane (DABCO, Sigma) 100 mg/ml. For microscopy, acquisition was done with an exposure time of 50 ms for Trans-DIC, 20 ms for DAPI 405, 200 ms for FITC and 600 ms for Tx2. Z stacking was used for imaging DAPI, FITC and Tx2 with a step of 0,3µm, ±3 µm around the acquisition point.

### Preparation of cytosol and polysome fractions

Polysome profiling was performed as described previously (26). Briefly, cells treated or not with CisPt were incubated at 37 °C with 100 μg/mL cycloheximide in fresh medium for 5 min. Cells were then washed, scraped into ice-cold PBS supplemented with 100 μg/mL cycloheximide, centrifuged at 3000 r.p.m. for 5 min. The cell pellets were resuspended into 400 μL of LSB buffer (20 mM Tris, pH 7.4, 100 mM NaCl, 3 mM MgCl2, 0.5 M sucrose, 1 mM DTT, 100 U/mL RNasin and 100 μg/mL cycloheximide). After homogenization, 400 μL LSB buffer supplemented with 0.2 % Triton X-100 and 0.25 M sucrose was added. Samples were stayed on ice for 30 min and centrifuged at 12,000 g for 15 min at 4 °C to pellet nuclei. The supernatant (cytosolic extract) was adjusted to 5 M NaCl and 1 M MgCl2. The lysates were then loaded onto a 5-50% sucrose density gradient and centrifuged in an SW41 Ti rotor (Beckman) at 36,000 rpm for 2 h at 4 °C. Fractions were monitored and collected using a gradient fractionation system (Isco). RNA was extracted from the four heaviest polysomal fractions (pooled) for 3’-seq analysis or from each fraction for RT-qPCR analysis.

### RNA extraction

RNA from whole cells or fractionated cell lysates were extracted with TRIzol Reagent (TRIzol-LS for polysomes; ThermoFischer Scientific) according to the manufacturer’s instructions, and 1 µl of GlycoBlue (ThermoFisher Scientific) was added for RNA precipitation. RNA from whole cells was treated with DNase I (TURBO DNA-free, ThermoFisher Scientific). RNA samples were quantified using a Nanodrop 2000 spectrophotometer (ThermoFischer Scientific). For sequencing, RNA samples were analyzed using an RNA 2100 Bioanalyzer (Agilent).

### RT-(q)PCR

Reverse transcription was performed on RNA using SuperScript III Reverse Transcriptase (ThermoFisher Scientific) and random primers. PCR was performed using GoTaq Flexi DNA Polymerase (Promega), and PCR products were migrated on agarose gels. Quantitative PCR (qPCR) was performed using Power SYBR Green PCR Master Mix (ThermoFisher Scientific) on a CFX96 Real-Time PCR Detection System (BioRad).

### 3’-seq experiments and bioinformatic analysis

5 μg of RNA were used for poly(A)+ RNA purification with Dynabeads mRNA DIRECT Micro kit (ThermoFisher Scientific). Poly(A)+ RNA was fragmented at 70°C for 5 min in RNA Fragmentation Reagents (ThermoFisher Scientific). After ethanol precipitation, RNA was reverse transcribed using anchored oligo-dT and SMARTScribe RT enzyme (Clontech) allowing to add adapter sequences to the 5’ and 3’ ends of RNA fragments. Libraries were amplified by 16 cycles of PCR with GoTaq Flexi DNA Polymerase (Promega), purified using Agencourt AMPure XP (Beckman Coulter) magnetic beads, and quantified with Quant-iT Picogreen dsDNA kit (ThermoFisher Scientific). A size selection step was done using SPRIselect (Beckman Coulter) magnetic beads to obtain fragments of 150–300 bp. Purified libraries were controlled by capillary electrophoresis (LabChip - Perkin) and quantitated by qPCR (KAPA Library Quantification Kits Illumina Platforms - Roche). Pooled libraries (12 pM) with 30% of phiX were subjected to single-end, 50 bp sequencing using the HiSeq 2500 machine (Illumina). Read 1 was read with primer HP6 (Illumina) with 3 dark cycles (first 3 bases of read 1 were not read). Index i7 (6 pb barcode) was read with primer HP8 (Illumina).

Raw reads were trimmed in their 5’ and 3’ ends to remove uninformative nucleotides due to primer sequences, nucleotides added by SMART Scribe RT enzyme and polyA tail of mRNAs. Trimmed reads of 25 bp or more were aligned on the human reference genome (hg19) using Bowtie2 (version 2.2.5) (27). Only reads with a mapping quality score (MAPQ) of 20 or more were retained (Samtools version 1.1) for downstream analysis. Reads were then clustered along the genome using Bedtools (version 2.17.0) (28), allowing a maximum distance of 170 bp (max library fragment length (300bp) – min fragment length (60bp) – read length (50bp) – oligodT length (25bp)) and a minimum number of 5 reads per peak. Peaks with a stretch of 6 consecutive As (or 8 As out of 9 nucleotides) within 150 bp downstream were filtered out, as they are likely due to internal priming of oligo-dT. Overlapping peaks from all analyzed samples were merged to define a common set of genomic windows corresponding to polyA sites. To annotate peak location within genes, gene coordinates were obtained on the basis of overlapping Refseq transcripts with the same gene symbol. Peaks overlapping any intronic region of a gene were classified as intronic polyA (IPA) peaks. Peaks overlapping the last exon of a gene were classified as LE peaks. Differential analyses between two conditions were done using three independent biological replicates per condition. To compare the regulation of each IPA to the regulation of the gene’s last exon (taken as the sum of the peaks in this exon), we used DESeq2 (version 1.4.5) (29) and the following statistical model:

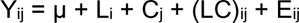

where Y_ij_ is the normalized counts of peak i in biological condition j, μ is the mean, L_i_ is the peak localization (IPA or LE), C_j_ is the biological condition, (LC)_ij_ is the interaction between peak localization and biological condition, and E_ij_ is the residual. P-values and adjusted P-values (Benjamini-Hochberg) were calculated. For H358 cells, data using a false discovery rate (FDR) of 10% are shown. For A549 cells, data with p < 0.05 are shown, to avoid under-estimations of cross-comparisons between lists. The complete bioinformatics pipeline (3’-SMART package) described above can be freely downloaded at GitHub (https://github.com/InstitutCurie/3-SMART) and can be run through a configuration file and a simple command line. Annotated polyadenylation sites were retrieved from the Polya_DB 3 and PolyASite 2.0 databases (30, 31).

### Long-read RNA-seq (Iso-seq) and bioinformatic analysis

A starting amount of 1 µg was used for long read library preparation. The first step of cDNA synthesis was done using the SMARTer PCR cDNA Synthesis Kit (Takara). Then, cDNA was amplified by 14 PCR cycles and divided in two fractions to be purified with different AMPure PB beads (Pacific Biosciences) ratio. In order to enrich for larger transcripts, a ratio of 0.4x was applied to a fraction (corresponding to 5 out of 8 of the initial volume), while the other fraction was purified with 1x ratio. For each sample, both fractions were quantified with Qubit dsDNA high-sensitivity kit (ThermoFisher Scientific), qualified by capillary electrophoresis (Bioanalyzer High Sensitivity DNA kit - Agilent), pooled equimolarly, and subjected to the SMRTbell Template Prep Kit 1.0 (Pacific Biosciences). The first DNA damage and end repair steps were applied according to the manufacturers’ recommendations. The following ligation was extended to 16 hours to improve efficiency. The final exonuclease treatment was applied to digest unwanted molecules. The SMRTbells were quantified with Qubit dsDNA high-sensitivity kit (ThermoFisher Scientific) and qualified by capillary electrophoresis as above. Complexes were prepared following the manufacturer’s recommendations using the Sequel Binding Kit 3.0 and the sequencing Primer v3 (Pacific Biosciences). Each sample was sequenced on 4 SMRTcells 1M, on the Sequel I system (Pacific Biosciences) using a loading molarity of 5 pM. For each SMRTcell, 10 hours of movie, 2 hours of immobilization, and 4 hours of pre-extension were set up. For bioinformatic analysis, subreads coming from PacBio SEQUEL SMRTcells were processed via the SMRTLink’s IsoSeq v3.0 pipeline with default parameters (https://github.com/PacificBiosciences/IsoSeq/blob/master/isoseq-clustering.md). This pipeline is composed of the following 6 steps: Circular Consensus Sequence (CCS) calling, primer removal/demultiplexing, refining, merging, clustering and polishing. Resulting CCS reads and HQ transcripts were mapped with minimap2 (v2.17-r941) against hg19 genome with the following parameters: -ax splice -uf --secondary=no -C5 -O6,24 -B4. Then, CCS reads were classified into ‘last exon (LE)’ and ‘intronic polyadenylation (IPA)’ isoforms, depending on whether the 3’-end of the read was located in the last exon of a gene or within a gene but upstream of the last exon.

### Total-RNA-seq and analysis of transcription processivity

For total-RNA-seq, total RNA from whole H358 cells treated or not with cisplatin (2 biological replicates of each condition) was subjected to DNAse I treatment. 500 ng of good quality RNA (RIN > 9) were used for Illumina compatible library preparation using the TruSeq Stranded total RNA protocol allowing to take into account strand information. A first step of ribosomal RNA depletion was performed using the RiboZero Gold kit (Illumina). After fragmentation, cDNA synthesis was performed and resulting fragments were used for dA-tailing followed by ligation of TruSeq indexed adapters. PCR amplification was finally achieved to generate the final barcoded cDNA libraries. Libraries were equimolarly pooled and subjected to qPCR quantification using the KAPA library quantification kit (Roche). Sequencing was carried out on the NovaSeq 6000 instrument from Illumina based on a 2*100 cycle mode (paired-end reads, 100 bases) using a S1 flow cell in order to obtain around 100 million clusters (200 million raw paired-end reads) per sample. Fastq files were generated from raw sequencing data using bcl2fastq where demultiplexing was performed according to barcodes. For each gene, the number of reads in the last intron was divided by the number of reads in the first intron. Genes with at least 20% reduction of this ratio in cisplatin *versus* vehicle conditions (p < 0.05) were considered to have their transcription processivity inhibited. A similar method was used to analyze intronic probes in exon-arrays.

### Other bioinformatic and statistical analyses

Intersection between lists of 3’-seq peaks were done using Bedtools (version 2.17.0) (28) allowing a gap of 300 bp (Fig. 2C). Functional gene annotation analyses were done using the DAVID software (32, 33), using the human genome as a reference. Genomic coordinates of sORFs were obtained from the sORFs.org (using a FLOSS classification of ‘good’) and OpenProt databases (34, 35) and were crossed with the genomic coordinates of cisplatin-regulated 5’UTR-IPA (from gene start to IPA site). For each experimental analysis, at least three independent experiments were performed. In bar charts, error bars represent the standard error of the mean (SEM) that is the standard deviation divided by the square root of sample number. Student’s paired t-tests were considered significant if p < 0.05.

### CRISPR-Cas9 edited clones

Single guide RNAs (sgRNAs) and a donor block oligonucleotide (for the mATG design) were designed using Integrated DNA technologies [IDT] design tool and those with the best on and off-target scores were selected (Table S3). PAM sequences were mutated on the donor block oligo to avoid recutting. For cell electroporation, 3.33 µg of Cas9 HiFi (IDT) in 0.5 µL of buffer R (Neon transfection Kit^TM^, Thermofisher) was mixed in a final volume of 1 µL with 22 picomol of sgRNA and incubated 10 to 20 min at room temperature. 10^5^ A549 cells were resuspended in 9 µL of Buffer R and added. 21.9 picomol of electroporation enhancer (IDT) and 22 picomol of donor block oligo (in the case of mATG) were added to the mix for a final volume of 11.5 µL. Electroporation was performed on a Neon transfector (ThermoFisher) at 1400 V with 4 pulses of 15 ms using Neon transfection kit 10 µL. Cells were seeded into 96 well plates. For mATG, media contained HDR enhancer V2 (IDT) at 1.34 µM and were changed 1 day after transfection. For cell cloning, 100 µL of cell suspension at 0.8 cell per 100 µL of medium (half fresh medium, half cell-conditioned medium filtered at 22 µm) were seeded in 96 well plates. For genomic DNA extraction, 25 µL of a trypsinized confluent cell plate was mixed with 25 µL of lysis buffer (20 mM Tris HCl pH 8.3, 100 mM KCl, 5 mM MgCl2, 1% NP-40, 1% Tween-20, 1:50 proteinase K). After 10 min at 65°C and 10 min at 95°C, high fidelity PCR was performed using Phusion enzyme in 20 µL final volume. Then, PCR products underwent either T7E1 enzymatic assay, restriction enzyme assay, or DNA cleaning for sequencing. For T7E1 enzymatic assay, 5 µL of the PCR product was dehybridized for 10 min at 95°C, then rehybridized by decreasing temperature to 85°C at 2°C/sec and to 25°C at 0.3°C/sec, then 1 U of T7 endonuclease 1 (New England Biolabs) and NEB Buffer 2 were added, samples were incubated for 1 hour at 37°C and analysed by electrophoresis on 1.2% agarose gel.

## RESULTS

### Cisplatin upregulates the IPA:LE isoform ratio in many genes

To analyze the impact of cisplatin on IPA isoform regulation, total RNA from the H358 human NSCLC cell line treated for 24 hours with either cisplatin or vehicle were analyzed by 3’-seq, that relies on the sequencing of regions preceding the 3’-terminal polyA tail of transcripts (36). Because 3’-seq analysis can be flawed by internal priming of the oligo(dT) at genomic stretches of A’s, peaks ending near a genomic stretch of A’s were discarded. We also verified that most of the selected peaks contained a potential polyadenylation signal (AAUAAA-like motif; Fig. S1A). For each peak overlapping an annotated intron, we analyzed the effect of cisplatin on its expression relative to the last exon (LE) of the cognate gene. With a false discovery rate (FDR) of 10%, 2963 intronic peaks in 1987 genes were regulated by cisplatin at 24 hours of treatment (Fig. 1A, ‘all’; Table S4). The vast majority of them (89%) were detected at a level of at least 5% of the LE of the gene (Fig. 1A, ‘abundant’). Because internal priming artefacts can be difficult to filter out bioinformatically, we also crossed our list of regulated intronic peaks with lists of already published polyA sites identified in normal tissues (30, 31). Among the 2,963 regulated intronic peaks, 1197 (40%) matched an annotated polyA site (Fig. 1A, ‘annotated’). Also, we found a similar percentage of annotated polyA sites in our subsequent 3’-seq analyses on cytosolic fractions (see below), which are unlikely to be subject to internal priming in introns. These data suggest that many of the cisplatin-regulated intronic peaks that we identified may correspond to genuine polyA sites, that were not annotated in normal tissues. We therefore decided to keep all regulated peaks for subsequent analyses.

**Figure 1:**
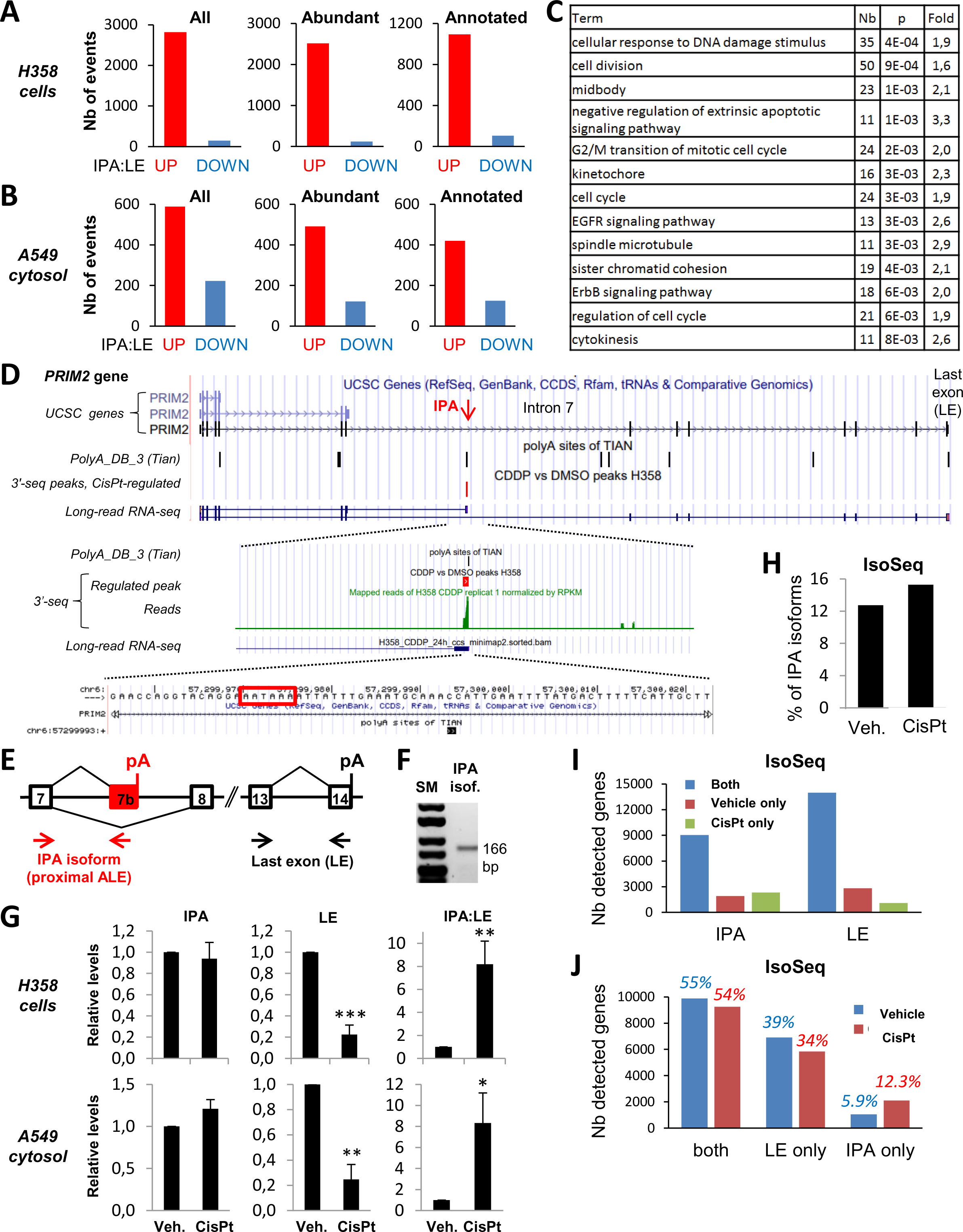
Cisplatin upregulates the IPA:LE isoform ratio in many genes. **A**, 3’-seq analysis of IPA:LE isoform ratio regulation by cisplatin in whole H358 cells treated with cisplatin or vehicle for 24 hours. **B**, Similar 3’-seq analysis in A549 cells (cytosol) treated with cisplatin or vehicle for 16 hours. **C**, Gene Ontology terms enriched in genes with IPA:LE regulation by cisplatin in H358 cells. Analysis with the DAVID software; the fold enrichment is indicated; only a subset of enriched functions are shown. **D**, Visualization of 3’-seq and long-read RNA-seq data for the *PRIM2* gene in the UCSC genome browser. The sequence around the regulated IPA site is shown and the polyadenylation signal is framed. **E**, Schematics of the *PRIM2* gene and PCR primers. **F**, RT-PCR detection of the PRIM2 IPA isoform. SM, size markers. **G**, RT-qPCR analysis of PRIM2 isoforms in H358 and A549 cells treated for 24 or 16 hours, respectively, with either cisplatin (CisPt) or vehicle (Veh.). **H-J**, Analysis of IPA and LE transcripts by long-read RNA-seq (Iso-Seq, PacBio) in H358 cells treated with cisplatin or vehicle for 24 hours. All detected transcripts were classified into IPA and LE transcripts. **H**, Percentage of IPA transcripts among detected transcripts. **I-J**, Number of genes detected at the level of IPA or LE in the presence or absence of cisplatin.

Strikingly, about 95% of IPA regulation events by cisplatin corresponded to an upregulation of the IPA:LE isoform ratio (Fig. 1A). This was in sharp contrast with the previously described effect of doxorubicin, which repressed the IPA:LE ratio in the majority of cases (7, 11), but was reminiscent of the effect of ultraviolet C (UV-C) irradiation (8, 9). Out of 84 IPA isoforms that were found to be upregulated by UV-C by RNA-seq in a previous study (8), 34 (40%) were also upregulated by cisplatin (Fig. S1B).

Likewise, 3’-seq analysis on another human NSCLC cell line (A549 cells) treated for 16 hours with cisplatin or with vehicle showed large-scale upregulation of IPA:LE isoform ratio (Fig. 1B and Table S5). 52% of the upregulation events in A549 cells were also found in H358 cells (Fig. S1C). Thus, large-scale up-regulation of IPA:LE isoform ratio is a robust effect of cisplatin in two different NSCLC cell lines. Analysis of gene functional annotation indicated that genes with cisplatin upregulation of the IPA:LE ratio were enriched in functions related to the DNA damage response, cell cycle and cell death (Fig. 1C), which are known to be impacted by cisplatin. Indeed, cisplatin induced cell cycle arrest and cell death in a time and dose-dependent manner (Fig. S1D).

An example of regulated gene is *PRIM2* (DNA Primase Subunit 2), which encodes a regulatory subunit of DNA primase, a DNA replication enzyme. In *PRIM2* intron 7, our 3’-seq data identified an IPA site, which is annotated in PolyA_DB_3 (chr6:57299993:+) and the expression of which was increased by cisplatin relative to the LE of the gene (exon 14; Fig. 1D). Although no RefSeq transcript matched this IPA site, long-read RNA-seq data that we generated in cisplatin-treated H358 cells showed the existence of a spliced mRNA containing exon 7 and an ALE within intron 7, matching the regulated IPA site (Fig. 1D). This IPA isoform was confirmed by RT-PCR analysis using primers located in exon 7 and the ALE (Fig. 1E-F). RT-qPCR analysis in both H358 and A549 cells validated the upregulation by cisplatin of the IPA:LE ratio of *PRIM2*, which was mainly due to a decrease of the LE isoform, while IPA isoform levels were maintained (Fig. 1G). This regulation by cisplatin was dose-dependent (Fig. S1E), led to decreased PRIM2 protein levels (Fig. S1F), and preceded cell death induction (Fig. S1D-E). The upregulation of IPA:LE isoform ratio by cisplatin was validated for additional genes by RT-PCR (Fig. S1G). In some genes (*ZFC3H1*, *HERC4*), the IPA isoform was readily increased by cisplatin, but in most cases, the regulation was merely due to a decrease of the LE isoform (Fig. S1G), as noted above for *PRIM2* (Fig. 1G). Finally, the IPA:LE ratio was not upregulated by cisplatin in the *HOMEZ* gene, used as a control (Fig. S1H).

To further assess the impact of cisplatin on IPA *versus* LE isoform regulation, we used single-molecule long-read RNA-seq (Iso-Seq) in H358 cells treated or not with cisplatin. We obtained more than one million transcripts in each condition (Fig. S1I). For each detected transcript, we determined whether it terminated within the LE of a gene (LE transcript) or upstream of the LE (IPA transcript). Cisplatin globally increased the proportion of IPA compared to LE transcripts by 20% (going from 12,7% to 15,3%; Fig. 1H); this represents a substantial increase when considering that the IPA:LE isoform ratio was upregulated in only 1875 genes in our 3’-seq data. Furthermore, there were 2.53 times less LE transcripts that were detected in cisplatin-treated cells only, when compared to vehicle-treated cells only; meanwhile, there were 1.22 times more IPA transcripts (Fig. 1I). Consistently, the fraction of genes detected at the IPA but not LE level was increased by 2.08-fold by cisplatin treatment, going from 5.9% to 12.3%; meanwhile, the fraction of genes detected at the LE but not IPA level was slightly decreased, going from 39% to 34% (1.15-fold decrease; Fig. 1J). Thus, both our 3’-seq and long-read RNA-seq data indicate that cisplatin globally favors the expression of IPA isoforms when compared to LE isoforms.

### A subset of IPA isoforms are depleted in heavy polysomes and terminate in the annotated 5’UTR part of genes

To gain insights into the potential function of IPA isoforms, and because little is known about IPA isoform translation on a genome-wide scale, we carried out 3’-seq analyses on the heavy polysomes fraction (HP, corresponding to actively translated mRNAs) of cisplatin- and vehicle-treated A549 cells. As in the case of cytosolic RNA that was analyzed in parallel (Fig. 1B), in the HP fraction cisplatin mostly upregulated the IPA:LE isoform ratio (Fig. 2A and Table S6). These data suggested that at least a subset of cisplatin-upregulated IPA isoforms are translated.

**Figure 2:**
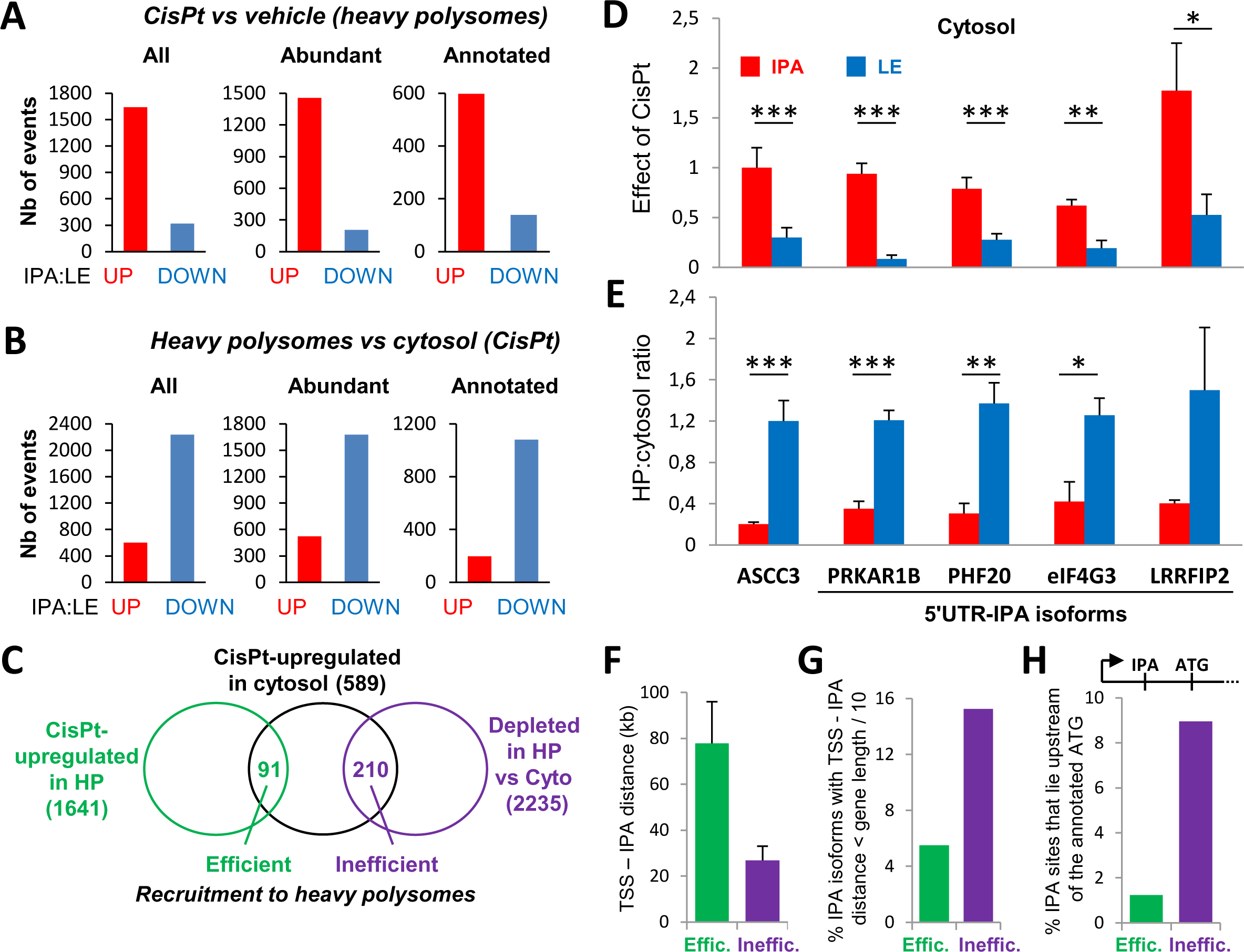
A subset of IPA isoforms are depleted in heavy polysomes and terminate in the annotated 5’UTR part of genes. **A**, 3’-seq analysis of IPA:LE isoform ratio regulation by cisplatin in heavy polysomes (HP) of A549 cells treated with cisplatin or vehicle for 16 hours. **B**, 3’-seq analysis of IPA:LE isoform ratio regulation in HP *versus* cytosol from cisplatin-treated A549 cells. **C**, Subsets of cisplatin-upregulated IPA isoforms are efficiently or inefficiently recruited to HP. Comparison between lists of IPA isoforms that are regulated relative to matched LE isoforms in the indicated conditions. **D-E**, RT-qPCR analysis of IPA and LE isoforms of the indicated genes, in the cytosol of A549 cells treated with cisplatin or vehicle (**D**) and in HP and cytosol of cisplatin-treated A549 cells (**E**). **F-H**, Comparison between the efficiently and inefficiently HP-recruited subsets of cisplatin-upregulated IPA isoforms identified in panel C. **F**, Distance from transcription start site (TSS) to IPA site. Median and SD. **G-H**, Percent of IPA isoforms that are located in the 5’ part of the gene (**G**) or upstream of the annotated translation initiation site (indicated as ATG) of the gene (**H**).

To compare the translation efficiency of IPA and LE isoforms, we then analyzed the relative abundance of IPA and matched LE isoforms in HPs *versus* cytosol. Performing this analysis on all IPA isoforms (not just those regulated by cisplatin), we observed that a subset of them exhibited a differential translation efficiency (more often lower) relative to the corresponding LE isoform (Fig. 2B and S2A; Tables S7 and S8). Thanks to these analyses, we identified a subset of cisplatin-upregulated IPA isoforms that were efficiently translated: they were upregulated by cisplatin (relative to matched LE isoforms) in both cytosol and HPs, and did not have a translation efficiency lower than the corresponding LE isoform (N=91; Fig. 2C and Table S9). These IPA isoforms likely encode protein isoforms with a truncated or distinct carboxy-terminal domain, compared to the protein encoded by the full-length mRNA (LE isoform), as previously shown for various genes (3).

These analyses also identified a subset of cisplatin-upregulated IPA isoforms that were inefficiently recruited to HPs: they were upregulated by cisplatin (relative to matched LE isoforms) in the cytosol but not in HPs, and had a lower HP:cytosol ratio than the corresponding LE isoform (N=210; Fig. 2C and Table S9). One of these 210 IPA isoforms is produced by the *ASCC3* (Activating Signal Cointegrator 1 Complex Subunit 3) gene and was previously reported to have a noncoding RNA function (8). In agreement with our 3’-seq data, RT-qPCR analyses showed that the *ASCC3* IPA isoform was upregulated (relative to the LE isoform) by cisplatin in the cytosol (Fig. 2D, left) and had a much lower HP:cytosol ratio than the LE isoform (Fig. 2E, left), which is consistent with its noncoding function.

When compared to the IPA isoforms that were efficiently translated (recruited to HPs), the inefficiently translated ones were on average 2.9 times shorter (Fig. 2F). In addition, they were enriched 2.8 times in the 5’ part of genes (Fig. 2G and S2B) and 7.3 times in the annotated 5’UTR part of genes (Fig. 2H). The latter configuration means that the IPA isoform terminates upstream of the annotated translation initiation site of the gene, which could explain the defect in recruitment to HPs. We thus coined such isoforms 5’UTR-IPA isoforms. (Of note, the *ASCC3* IPA isoform is not a 5’UTR-IPA isoform because it is produced from the coding region of the gene.)

For four genes with 5’UTR-IPA isoforms, namely *PHF20* (PHD Finger Protein 20), *PRKAR1B* (Protein Kinase cAMP-Dependent Type I Regulatory Subunit Beta), *EIF4G3* (Eukaryotic Translation Initiation Factor 4 Gamma 3) and *LRRFIP2* (LRR Binding FLII Interacting Protein 2), we validated by RT-qPCR that the 5’UTR-IPA isoform was upregulated (relative to the LE isoform) by cisplatin in the cytosol and had a much lower HP:cytosol ratio than the LE isoform (Fig. 2D-E). In other words, for these genes, the IPA:LE isoform ratio was upregulated by cisplatin in the cytosol, and was much lower in HPs than in the cytosol, in agreement with our 3’-seq data. Altogether, our 3’-seq data identify a set of IPA isoforms, especially 5’UTR-IPA isoforms, that are upregulated (relative to the LE isoform) by cisplatin in the cytosol, but are depleted in HPs when compared to cytosol.

### The 5’UTR-IPA isoforms of the *PHF20* and *PRKAR1B* genes impact cisplatin sensitivity

To determine whether the *PHF20* and *PRKAR1B* 5’UTR-IPA isoforms may play a functional role in cisplatin response, we designed two independent siRNAs that depleted each IPA isoform without affecting the corresponding LE isoform (Fig. 3A). In A549 cells transfected with a control siRNA, cisplatin decreased cell growth by about 60% (data not shown). siRNA-mediated depletion of both IPA isoforms had no robust effect on cell growth in the absence of cisplatin (Fig. 3B), but reproducibly increased cell survival to cisplatin, that is, cell growth in the presence of cisplatin normalized to cell growth without drug (Fig. 3C). These data indicate that the *PHF20* and *PRKAR1B* 5’UTR-IPA isoforms, whose expression is maintained following cisplatin treatment (Fig. 2D), inhibit cell survival to cisplatin. Because cisplatin decreases the expression levels of the LE isoforms of these genes (Fig. 2D), we also determined the effects of depleting these isoforms on cell growth. For both genes, two independent siRNAs that depleted the LE but not the IPA isoform led to decreased cell growth in both the absence and presence of cisplatin (Fig. 3D-E). Thus, for both genes, our data suggest that in response to cisplatin, cell growth is inhibited both by the 5’UTR-IPA isoform (whose expression is maintained) and by the decreased expression of the canonical LE isoform (Fig. 3F).

**Figure 3:**
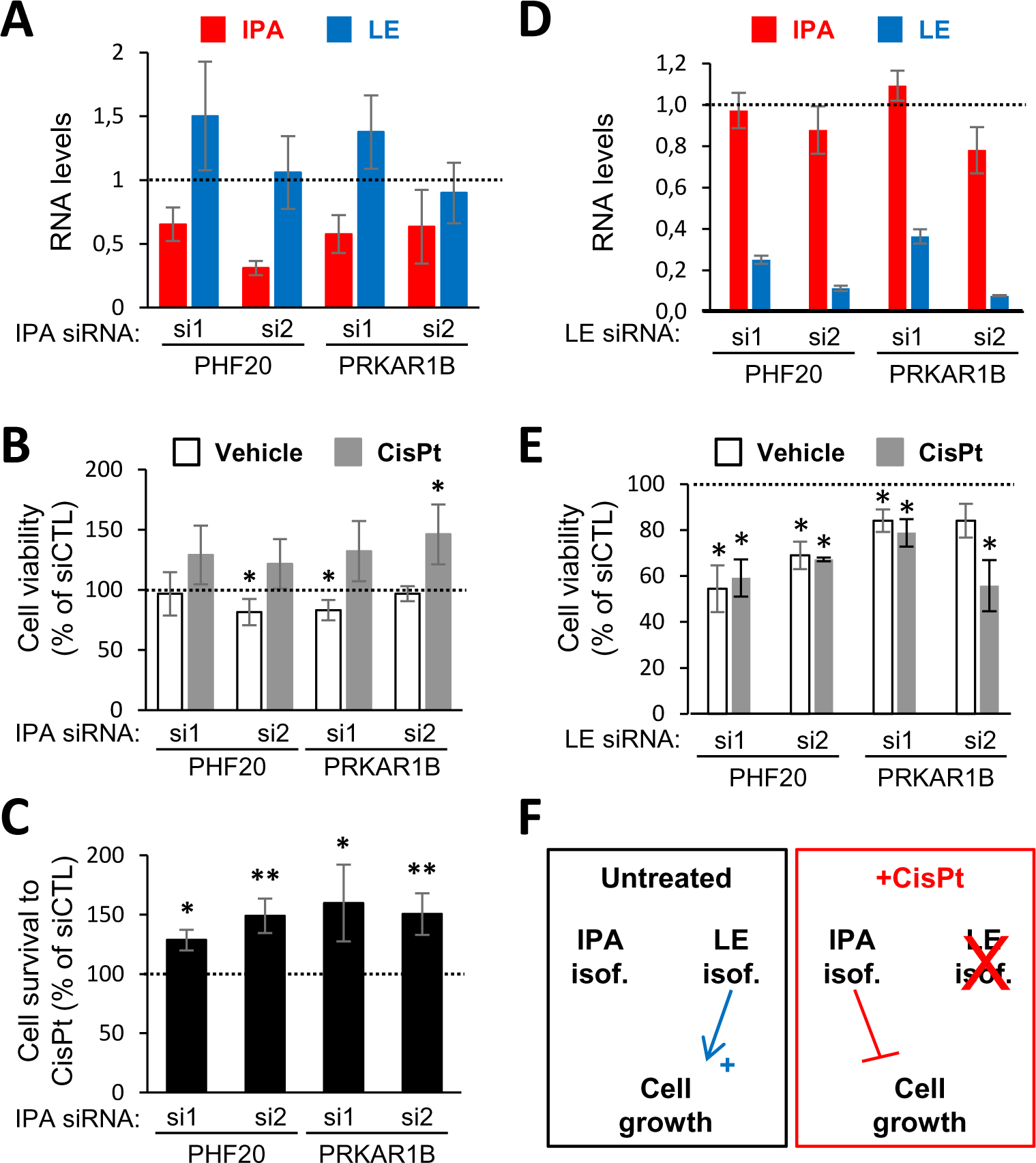
The *PHF20* and *PRKAR1B* 5’UTR-IPA isoforms impact cisplatin sensitivity. Effects of transfected siRNAs targeting either the IPA or LE isoform of *PHF20* or *PRKAR1B* in A549 cells. Two independent siRNAs were used for each isoform. Data in the presence of a negative control siRNA (siCTL) were set to 100% or 1. **A** and **D**, RT-qPCR analysis of IPA and LE isoforms abundance in vehicle-treated cells. **B**, **C** and **E**, WST1 assay analysis of A549 cell viability in the presence or absence of cisplatin for 24 hours. In C, cell viability with cisplatin was normalized to cell viability without cisplatin. **F**, Model showing the impact of the IPA and LE isoforms of *PHF20* and *PRKAR1B* on cell growth in the absence and presence of cisplatin.

### The 5’UTR-IPA isoforms of the *PHF20* and *PRKAR1B* genes are translated into microproteins

Because the cytosolic IPA isoforms of *PHF20* and *PRKAR1B* are depleted in HPs (Fig. 2E) and are generated from the annotated 5’UTR part of genes, we reasoned that their depletion phenotype (Fig. 3) could be due either to a noncoding function or to the production of microproteins encoded by sORFs (see introduction) (18, 19). To determine whether 5’UTR-IPA isoforms may contain sORFs, we first determined their exact exon content. The *PHF20* and *PRKAR1B* 5’UTR-IPA isoforms are not annotated. Our long-read RNA-seq data indicated that the *PHF20* 5’UTR-IPA isoform starts in the first exon of the gene, like the full-length *PHF20* mRNA (Fig. 4A). In the 5’UTR-IPA isoform, exon 1 is spliced to a non-annotated exon that is located within the first intron of the full-length transcript and that is supported by both our long-read RNA-seq and total-RNA-seq data (that is, RNA-seq on total RNA depleted of ribosomal RNA). This exon ends with a polyA site, that is also supported by our 3’-seq data (Fig. 4A). The *PRKAR1B* gene has two alternative first exons, both of which can be used to produce the full-length *PRKAR1B* mRNAs. In this gene, our long-read RNA-seq data revealed two 5’UTR-IPA isoforms that contain either one of the two alternative first exons, which are then spliced to a non-annotated exon (Fig. 4B). This novel exon is also supported by our total-RNA-seq data and ends with a polyA site supported by our 3’-seq data. Thus, in both genes, the 5’UTR-IPA isoforms contain an ALE exon located within the annotated first intron of the gene.

**Figure 4:**
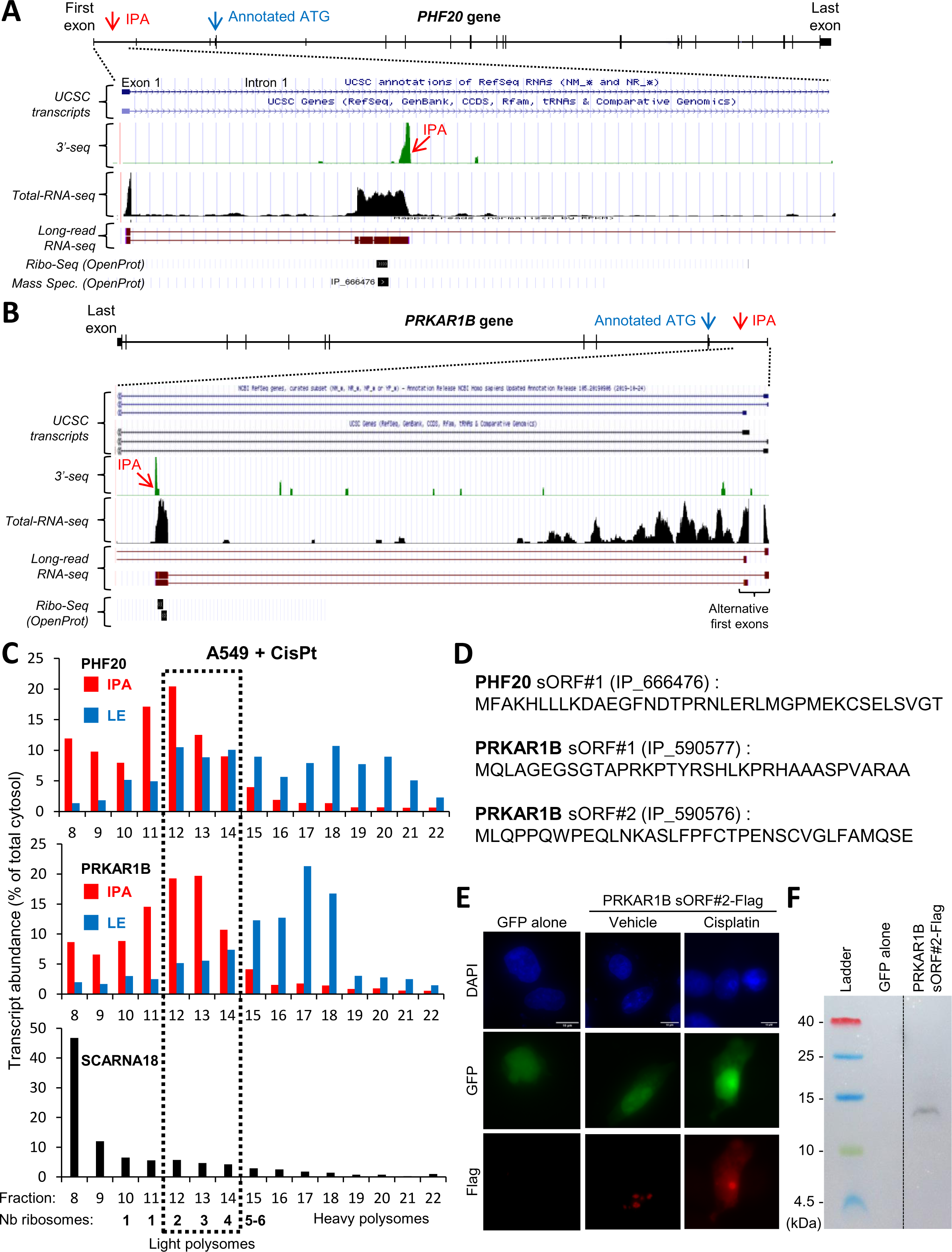
The *PHF20* and *PRKAR1B* 5’UTR-IPA isoforms are translated into microproteins. **A-B**, Visualization of our 3’-seq, total-RNA-seq and long-read RNA-seq data, as well as Ribo-seq and mass spectrometry data from OpenProt, for the 5’ part of the *PHF20* and *PRKAR1B* genes in the UCSC genome browser. The regulated IPA site and the annotated translation initiation site (ATG) are shown. **C**, RT-qPCR analysis of the *PHF20* and *PRKAR1B* IPA and LE isoforms and of of the *SCARNA18* lncRNA in sucrose gradient fractions prepared from the cytosol of cisplatin-treated A549 cells. The number of ribosomes in the different fractions is indicated. **D**, Sequence of the predicted microproteins encoded by sORFs in the *PHF20* and *PRKAR1B* 5’UTR-IPA isoforms. The OpenProt accession number of the sORFs is given. **E-F**, Analysis of cells transfected with expression plasmids encoding GFP and containing or not the *PRKAR1B* 5’UTR-IPA isoform with a Flag tag in frame with sORF#2. **E**, Anti-Flag immunofluorescence, GFP fluorescence, and DAPI staining (nuclei) in HeLa cells grown in the presence or absence of 10 µM Cisplatin for 16 hours. The bars represent 10 µm. **F**, Western blot analysis of HEK-293T cells.

We then searched for sORFs supported by Ribo-seq datasets and matching the *PHF20* and *PRKAR1B* 5’UTR-IPA isoforms. In the OpenProt database (34), one such sORF (hereafter referred to as *PHF20* sORF#1) was found for *PHF20* and two were found for *PRKAR1B* (sORF#1 and sORF#2) (Fig. 4A-B ‘Ribo-Seq’, Fig. S3A-B, and Table S10). For both genes, these sORFs are entirely contained within the ALE exon that defines the 5’UTR-IPA isoform. For *PHF20*, additional potential Ribo-seq-supported sORFs were found in the sORFs.org database (35) (Table S11). These data suggest that the *PHF20* and *PRKAR1B* 5’UTR-IPA isoforms are translated.

To further test whether the *PHF20* and *PRKAR1B* 5’UTR-IPA isoforms are translated in cisplatin-treated A549 cells, we analyzed by RT-qPCR the sucrose density gradient fractions that were obtained from the cytosol of these cells and that correspond to sub-, light- and heavy-polysomal fractions. For both genes, the 5’UTR-IPA isoforms were associated with lower-density fractions when compared to matched LE isoforms (Fig. 4C). The 5’UTR-IPA isoforms were almost absent from the heaviest polysomal fractions, while they were predominant in sub-polysomal fractions. Importantly, about 15% of each 5’UTR-IPA isoform was associated with a single full ribosome (corresponding to a recognized initiation codon), and about 45% with light polysomes (mainly 2 or 3 ribosomes; Fig. 4C). Similar results were obtained in vehicle-treated cells (Fig. S3D). As controls, the lncRNA *SCARNA18* was scarcely found in light polysomes (Fig. 4C, bottom), while the IPA isoform of *PATZ1* was abundant in HPs in agreement with our 3’-seq data (Fig. S3C and Table S7). These data indicate that the *PHF20* and *PRKAR1B* 5’UTR-IPA isoforms are enriched in light polysomes. Altogether, these polysome profiling data and the above-mentioned Ribo-seq data indicate that the *PHF20* and *PRKAR1B* 5’UTR-IPA isoforms are translated. Their association with light, rather than heavy polysomes, is consistent with the small size of their ORFs (coding 35 to 38 amino acids), given that a ribosome covers about 30 nucleotides.

Then, we investigated whether microproteins that are predicted to be produced by sORFs in the *PHF20* and *PRKAR1B* 5’UTR-IPA isoforms (Fig. 4D) could indeed be detected. In mass spectrometry datasets available in OpenProt (34), we found peptide detection evidence for the microprotein corresponding to sORF#1 of *PHF20* (Fig. 4A, ‘Mass Spec’) but no such evidence for the sORFs of *PRKAR1B*. To further test microprotein production, we cloned *PHF20* sORF#1, *PRKAR1B* sORF#1 and *PRKAR1B* sORF#2 in expression plasmids, with a carboxy-terminal Flag tag, the ALE sequence upstream of the sORF (the sequence downstream of the sORF was omitted), and a separately encoded GFP. Following transfection into HeLa cells, immunofluorescence with an anti-Flag antibody detected no signal in the case of a GFP-only plasmid (without Flag) but a strong signal in the case of *PRKAR1B* sORF#2 (Fig. 4E). In the absence of cisplatin, this signal was exclusively nuclear and mainly located in a few large nuclear areas (Fig. 4E, bottom, ‘vehicle’). However, following cisplatin treatment, the signal became diffuse throughout the nucleus and cytoplasm (Fig. 4E, bottom, ‘cisplatin’). We could not detect signal for *PRKAR1B* sORF#1 and *PHF20* sORF#1 (data not shown). The microprotein encoded by *PRKAR1B* sORF#2 was also detected by Western blot analysis on transfected HEK-293T cells (Fig. 4F). Thus, multiple approaches (*i.e.*, polysome profiling, Ribo-Seq, mass spectrometry, immunofluorescence, and/or Western blot) indicate that the 5’UTR-IPA isoforms of *PRKAR1B* and *PHF20* are translated and encode microproteins. Altogether, our data identify 5’UTR-IPA isoforms of *PHF20* and *PRKAR1B*, that are translated (Fig. 4) and inhibit cell survival to cisplatin (Fig. 3).

### The cisplatin survival phenotype of the *PRKAR1B* 5’UTR-IPA isoform is attributable to its sORF#2

To determine, whether the phenotype of the *PRKAR1B* 5’UTR-IPA isoform is due to its sORF#2, we mutated by CRISPR the initiation codon of this sORF in the endogenous *PRKAR1B* gene in A549 cells and we obtained 7 such homozygously mutated clones (Fig. 5A, mATG). We also generated 4 homozygous clones with CRISPR deletion of the IPA site (ΔIPA). As control cells, we used 8 unsuccessful CRISPR clones and parental A549 cells. We verified by RT-qPCR that the expression of the *PRKAR1B* 5’UTR-IPA isoform was compromised in ΔIPA (but not mATG) clones, when compared to control cells (Fig. 5B). In addition, Western blot analysis showed that the expression of the canonical PRKAR1B protein was not altered in ΔIPA and mATG clones (Fig. 5C). Finally, we looked at cell growth in the three groups of clones. In the absence of cisplatin, cell growth was slightly increased in ΔIPA but not mATG clones, when compared to control cells (Fig. 5D). In conditions where cell survival to cisplatin was about 60% in the control group, both the ΔIPA and mATG groups of clones survived significantly more to the drug, when compared to the control group (Fig. 5E). This increased survival to cisplatin, which was also observed with siRNAs targeting the *PRKAR1B* 5’UTR-IPA isoform (Fig. 3C), was found in the majority of both ΔIPA and mATG clones (Fig. 5E; a minority did not survive more than controls, possibly due to off-target mutations or compensatory mechanisms). These data indicate that both the depletion of the *PRKAR1B* 5’UTR-IPA isoform (by either siRNA or CRISPR) and the CRISPR mutation of its sORF#2 ATG lead to increased cell survival to cisplatin. Thus, the phenotype of the *PRKAR1B* 5’UTR-IPA isoform is due at least in part to its sORF#2.

**Figure 5:**
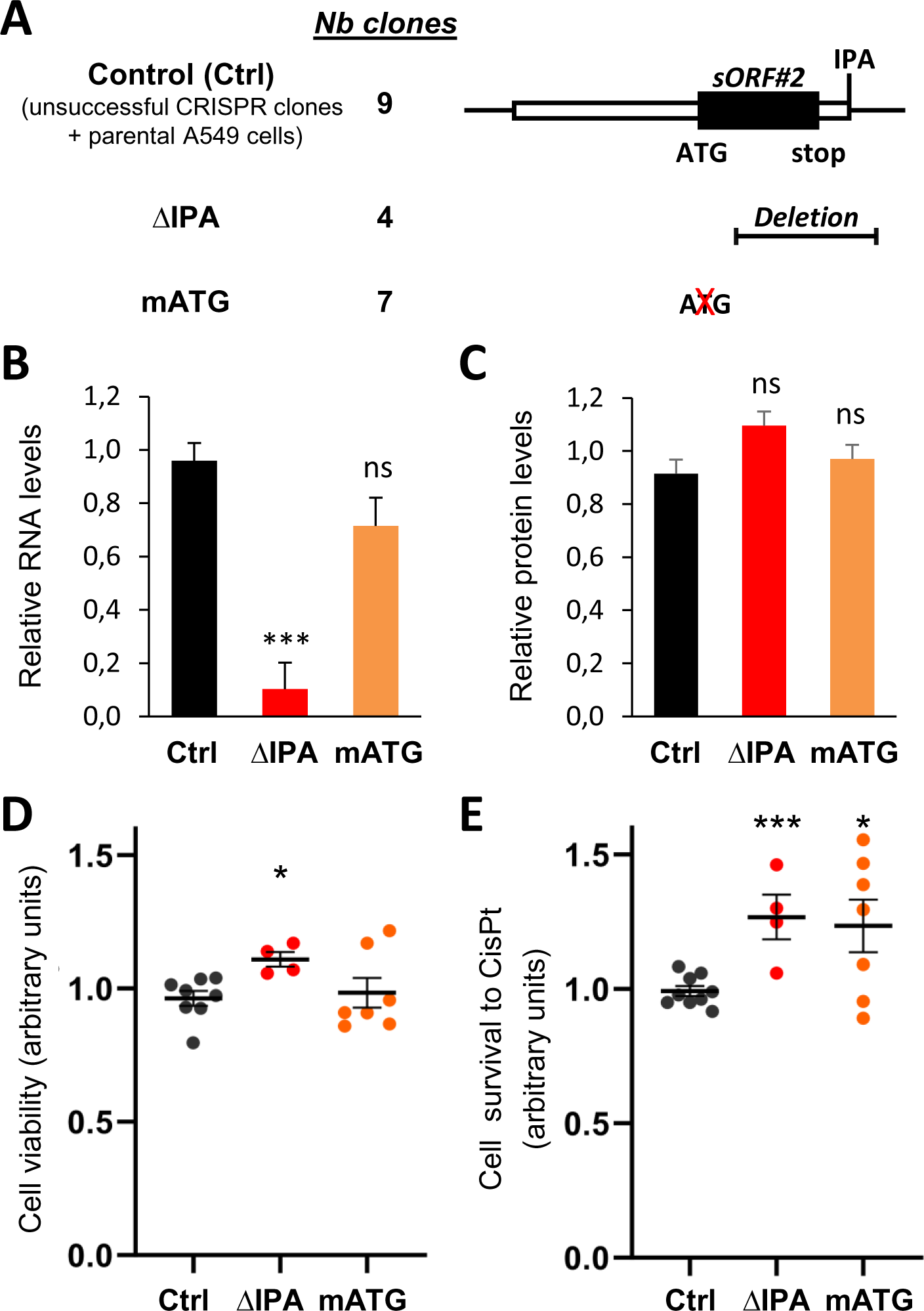
The cisplatin survival phenotype of the *PRKAR1B* 5’UTR-IPA isoform is attributable to its small ORF#2. **A**, Schematics of CRISPR-edited and control (Ctrl) cell lines. **B-E**, Analyses of the Ctrl, ΔIPA, and mATG cell lines. The graphs use arbitrary units. Statistical tests use Ctrl cells for comparison. **B**, RT-qPCR analysis of expression levels of the *PRKAR1B* IPA isoform, normalized to TBP mRNA levels. **C**, Western blot quantification of the PRKAR1B canonical protein, normalized to GAPDH protein levels. **D-E**, WST1 analysis of cell viability in the presence or absence of cisplatin for 24 hours. **D**, Vehicle-treated cells. **E**, Cell viability with cisplatin was normalized to cell viability without cisplatin.

### The concept of miP-5’UTR-IPA genes and their widespread regulation by cisplatin

Based on our data on the *PHF20* and *PRKAR1B* genes, we propose to define the novel paradigm of miP-5’UTR-IPA genes, where an IPA isoform is generated in the annotated 5’UTR part of a canonical protein-coding gene and encodes a microprotein (Fig. 6A). To further identify miP-5’UTR-IPA genes regulated by cisplatin, we first identified 290 genes with 5’UTR-IPA isoforms that were upregulated (relative to the last exon of genes) in response to cisplatin in either H358 or A549 cells (Table S12). We found that 156 of them had, between the gene 5’-end and the IPA site, an sORF supported by Ribo-Seq (135 cases) and/or mass spectrometry datasets (50 cases; Fig. 6B and Table S13). These data identify a set of 156 candidate genes with cisplatin-regulated miP-5’UTR-IPA. Looking at the annotated function of these 156 genes, we found them to be enriched in specific functions, most notably signal transduction and transcription regulation (Fig. 6C), as exemplified by *PRKAR1B* and *PHF20*, respectively.

**Figure 6:**
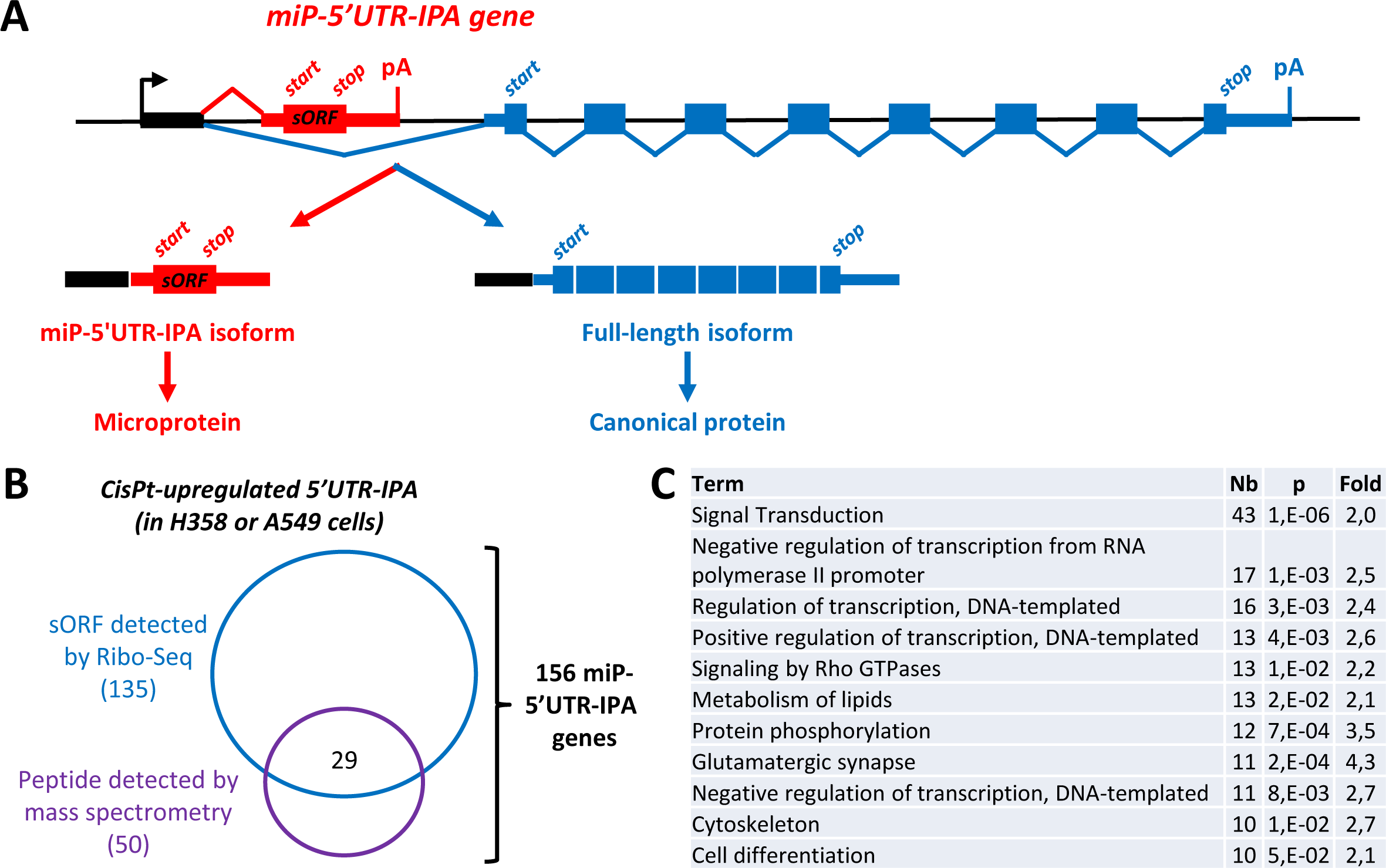
Genome-wide identification of putative miP-5’UTR-IPA isoforms upregulated by cisplatin relative to LE isoform. **A**, Model of the novel genetic paradigm of miP-5’UTR-IPA. **B**, Subsets of cisplatin-upregulated 5’UTR-IPA isoforms (from A549 and H358 cells) that contain sORFs supported by Ribo-Seq and/or mass spectrometry datasets. **C**, Gene Ontology terms enriched in the set of 156 miP-5’UTR-IPA candidate genes identified in B. Analysis with the DAVID software. Shown are functions that are enriched by at least 2-fold, with an adjusted p value below 0.05, and with at least 10 genes. Nb, number of genes. Fold, fold enrichment.

### Cisplatin effect on the IPA:LE ratio is mediated by decreased transcription processivity and depends on FANCD2 and senataxin

Finally, we investigated on the molecular mechanisms underlying IPA:LE isoform regulation in response to cisplatin. Based on our long-read RNA-seq data, the average length of detected mRNAs was 1.54 times smaller in cisplatin-treated cells than in vehicle-treated cells (1.3 *versus* 2.0 kb; Fig. 7A, left panel). The cisplatin-induced decrease in transcript length was even more pronounced (1.86-fold) when introns were taken into account (pre-mRNA length of 5.0 *versus* 9.3 kb; Fig. 7A, right panel). These data suggest that cisplatin favors the expression of shorter primary transcripts. In our 3’-seq data, genes with IPA:LE isoform ratio upregulation by cisplatin were about 3 times longer than genes with non-regulated IPA, indicating that the upregulation of IPA:LE isoform ratio by cisplatin is enriched in long genes (Fig. 7B). Altogether, our 3’-seq and long-read RNA-seq data indicate that cisplatin treatment favors the expression of IPA *versus* full-length (LE) isoforms in long genes. (Cisplatin also seemed to favor the expression of shorter genes, because pre-mRNA length of LE transcripts in cisplatin-treated cells was similar to that of IPA transcripts in vehicle-treated cells [Fig. 7A, right panel]; and the length of cisplatin-upregulated IPA transcripts was similar to the length of LE transcripts of non-regulated genes [Fig. 7B]).

**Figure 7:**
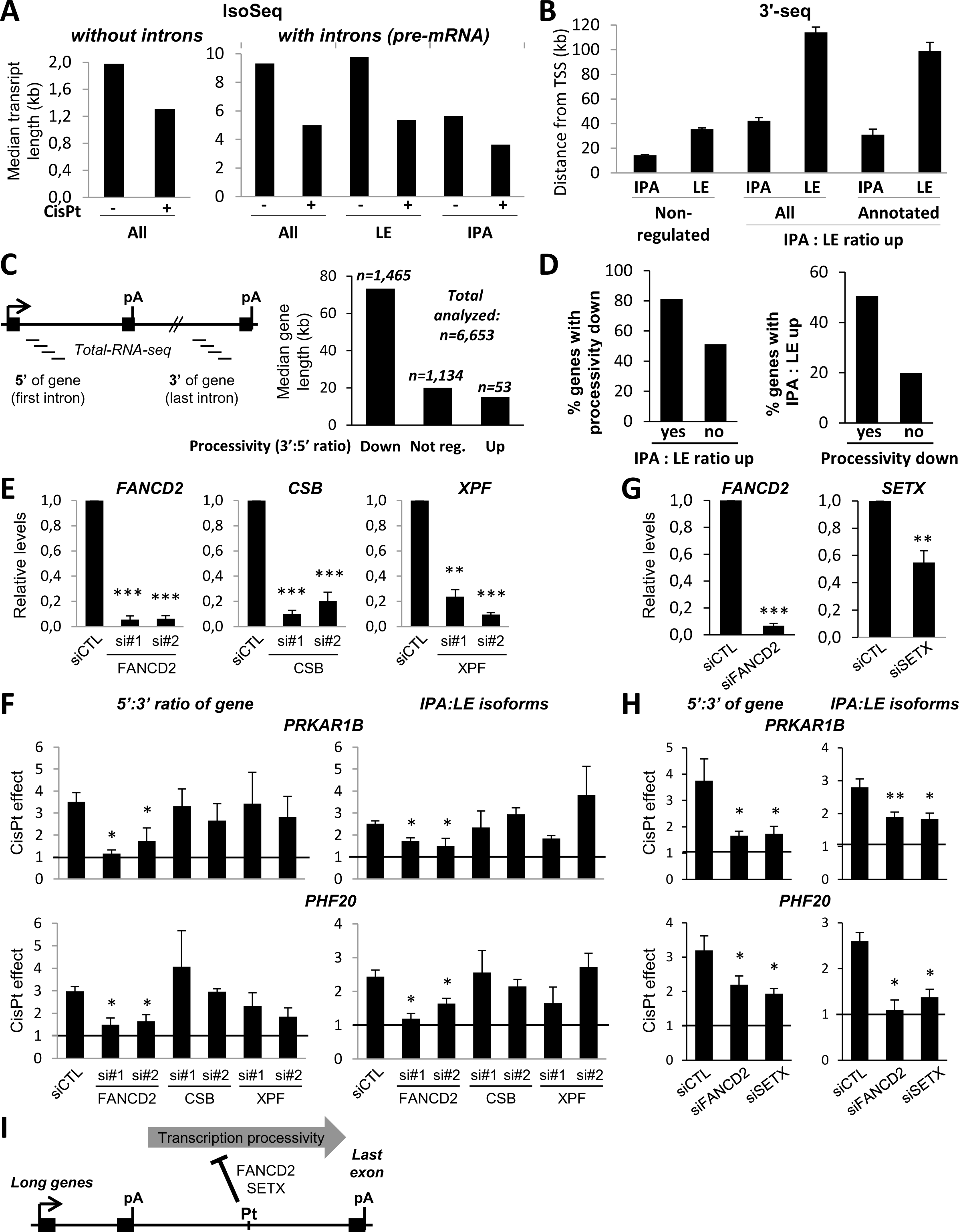
Cisplatin effect on the IPA:LE ratio is mediated by decreased transcription processivity and depends on FANCD2 and senataxin. **A**, Analysis of IPA and LE transcripts by long-read RNA-seq (Iso-Seq, from Fig. 1H) in H358 cells treated with cisplatin or vehicle for 24 hours. Median length of IPA, LE and overall transcripts detected in the presence or absence of cisplatin, either without or with introns taken into account. **B**, Distance (showing median and SD) from transcription start site (TSS) to polyA site of IPA and matched LE isoforms. Comparison between IPA isoforms that were either non-regulated or upregulated by cisplatin in 3’-seq analyses (from Fig. 1A). **C**, Analysis of transcription processivity by total-RNA-seq in H358 cells treated with cisplatin or vehicle for 24 hours. For each gene, processivity was assessed as the ratio of read number in the last *versus* first intron (3’:5’ ratio). Median length and number of genes, whose processivity was either up-, down- or non-regulated by cisplatin. **D**, Enrichment of IPA:LE isoform ratio upregulation in genes with processivity inhibition in response to cisplatin; and *vice versa*, enrichment of processivity inhibition in genes with IPA:LE isoform ratio upregulation. The regulation of IPA:LE isoform ratio and of processivity come from 3’-seq and total-RNA-seq analyses, respectively. **E-H**, A549 cells were transfected with siRNAs targeting the indicated factors or a control siRNA (siCTL) and were treated with cisplatin or vehicle for 8 hours. In G-H, *FANCD2* was depleted using a pool of the two siRNAs used in E-F. **E** and **G**, RT-qPCR analysis of the siRNA-targeted factors. **F** and **H**, RT-qPCR analysis of the fold-effects of cisplatin on the 5’:3’ ratio (that is, the inverse of processivity) and on the IPA:LE isoform ratio for the *PRKAR1B* and *PHF20* genes. **I**, Model of the molecular mechanism, by which cisplatin upregulates the ratio of (miP-5’UTR-)IPA to LE isoforms in long genes.

Because the cisplatin-induced upregulation of the IPA:LE isoform ratio was enriched in long genes and was accompanied by a decrease of LE isoform levels, both globally (Fig. 1I-J) and in many tested genes (Fig. 1G, 2D, and S1G), we reasoned that this cisplatin effect may be due to an inhibition of transcription processivity toward the 3’-end of long genes. Also consistent with this hypothesis, cisplatin was reported to stall RNA polymerase and inhibit transcription elongation in several genes or gene constructs (37–39). To test this hypothesis, we carried out total-RNA-seq (that is, RNA-seq on total RNA depleted of ribosomal RNA) on H358 cells treated or not with cisplatin, and we used the ratio of intronic reads at the 3’ *versus* 5’ part of genes (3’:5’ of gene) as a proxy for transcription processivity. Cisplatin inhibited the relative expression of the 3’ *versus* 5’ part of genes in 1,465 (22%) out of 6,653 analyzed genes (>20% decrease and p < 0.05) and this effect was enriched in long genes (Fig. 7C and Table S14), suggesting a decrease of transcription processivity. Genes exhibiting IPA:LE isoform ratio upregulation by cisplatin had more often their transcription processivity inhibited by the drug, when compared to genes with non-regulated IPAs (81% *versus* 51%; Fig. 7D, middle). Conversely, IPA:LE isoform ratio upregulation was more frequent in genes exhibiting processivity inhibition (50% *versus* 20%; Fig. 7D, right). Thus, the upregulation of IPA:LE isoform ratio and the downregulation of transcription processivity in response to cisplatin were correlated. By RT-qPCR using primers in introns at the 3’ *versus* 5’ part of genes, we verified that cisplatin treatment inhibited transcription processivity in 10 out of 13 tested genes with IPA:LE isoform ratio upregulation (Fig. S4A-B and see below). More detailed analyses on the *PRIM2* gene indicated that in response to cisplatin, pre-mRNA levels strongly decreased between the regulated IPA and the last intron (Fig. S4C); the dose-dependent effects on processivity and isoform ratio were correlated (Fig. 1H and S4D); and the processivity effect was seen at as early as 1 hour of treatment (Fig. S4E). Thus, the cisplatin-induced decrease of processivity likely explains the selective decrease of the LE but not IPA isoform levels and the increase in the IPA:LE isoform ratio.

We then investigated on factors that could explain the cisplatin-induced decrease of transcription processivity and increase of IPA:LE isoform ratio. Cisplatin-DNA crosslinks in transcribed genes trigger RNA polymerase II stalling and transcription-coupled nucleotide excision repair (TC-NER) mediated by CSB and XPF (XPF is also involved in other repair pathways) (40). In addition, interstrand DNA crosslinks induced by cisplatin are repaired by the Fanconi anemia pathway involving FANCD2 (40). Depletion of either CSB or XPF with siRNAs did not prevent cisplatin effects on processivity and IPA:LE isoform ratio in the *PRKAR1B* and *PHF20* genes and on *PRIM2* processivity (Fig. 7E-F and S4F), thus arguing against a role of TC-NER in these cisplatin effects. In contrast, siRNA depletion of FANCD2 with two independent siRNAs partially prevented cisplatin effects on processivity and IPA:LE ratio in the *PRKAR1B* and *PHF20* genes and on *PRIM2* processivity (Fig. 7E-F and S4F). These effects of FANCD2 are unlikely due to its canonical role in the Fanconi repair pathway, that occurs in S phase (40). FANCD2 was also shown to interact with R-loops (41–44), which are structures made of RNA:DNA hybrids (and a displaced DNA strand) and formed during transcription; and to colocalize with senataxin (SETX) (45), an R-loop helicase that promotes the use of some polyA sites (46). As in the case of FANCD2, siRNA depletion of SETX partially prevented cisplatin effects on processivity and IPA:LE ratio in the *PRKAR1B* and *PHF20* genes and on *PRIM2* processivity (Fig. 7G-H and S4G). Thus, our data suggest that the cisplatin-induced increase in the ratio of (miP-5’UTR-)IPA to LE isoforms in long genes is mediated, at least in part, by a decrease of transcription processivity, that is dependent on FANCD2 and SETX (Fig. 7I).

## DISCUSSION

IPA isoforms are widely regulated in various biological processes and are usually thought to encode protein isoforms (they are sometimes referred to as coding region-APA isoforms (4, 47)). Our findings that many cisplatin-regulated IPA isoforms are generated in the annotated 5’UTR part of genes and are translated into microproteins provide new insights into the translation outcome and function of IPA isoforms and reveal the novel paradigm of miP-5’UTR-IPA genes. In addition, our findings that cisplatin upregulates the IPA:LE isoform ratio in many genes provides new insights into the effects of this anticancer drug and suggest a role for miP-5’UTR-IPA isoform regulation in cancer cell response to cisplatin.

The almost systematic up- (as opposed to down-) regulation pattern of the IPA:LE isoform ratio in whole cells in response to cisplatin is distinct from the effects previously described for other genotoxic anticancer drugs, with doxorubicin downregulating the IPA:LE isoform ratio in most cases and camptothecin inducing both patterns in approximately equal proportions (7, 11). Our data strongly suggest a role for transcription processivity inhibition in IPA:LE isoform ratio upregulation by cisplatin. Previous analyses on gene constructs showed that cisplatin-DNA adducts induce RNA polymerase stalling and inhibit transcription elongation; several mechanisms are involved, including interstrand covalent bonds (crosslinks), RNA polymerase modifications, and reduced nucleosome mobility (37–39, 48, 49). Our analyses of intronic RNA levels by total-RNA-seq in thousands of endogenous genes and by RT-qPCR in several genes (Fig. 7 and S4A-E) suggest that cisplatin inhibits transcription processivity in long genes, which can directly explain the decrease of full-length mRNA (LE isoform) levels relative to IPA isoform levels. In future studies, analyses of nascent RNA could be used to more directly assess transcription dynamics, including elongation rate. Because cleavage/ polyadenylation is extensively coupled to transcription, reduced elongation rate near the IPA site could favor its cotranscriptional recognition, which in turn could favor intronic transcription termination. Upregulation of the IPA:LE isoform ratio through elongation inhibition in long genes was proposed for UV-C (8); consistently, many regulation events induced by UV-C were found in our cisplatin dataset (Fig. S1B; the limited dataset available for UV-C precludes the converse analysis).

Mechanistically, our data indicate that processivity inhibition and IPA:LE upregulation by cisplatin are dependent on FANCD2 and SETX (Fig. 7E-I and S4F-G). Both of these proteins are known to colocalize with each other (45) and to interact with R-loops (41–44), and the effects of SETX on cleavage/ polyadenylation are R-loop dependent (46). Consistent with this, we found that depletion of RNAse H1 (which selectively degrades RNA hybridized to DNA) prevented processivity inhibition by cisplatin in the *PRIM2* gene (Fig. S4H-I), but effects on other genes were more elusive (data not shown). Thus, the potential role of R-loops in IPA:LE regulation by cisplatin, FANCD2 and SETX remains to be investigated. Additional factors may contribute to IPA:LE upregulation by cisplatin, because IPA is regulated by many factors involved in cleavage/ polyadenylation, splicing, transcription elongation and termination, and epigenetic marks (1–4). For example, depletion or inhibition of CDK12 (a protein kinase that phosphorylates RNA polymerase II) inhibits transcription processivity and upregulates the IPA:LE isoform ratio preferentially in long genes (13, 50). Finally, IPA:LE isoform ratio upregulation in several genes upon UV-C irradiation was shown to be mediated by downregulation of the U1 snRNA, which as part of the U1 snRNP is a widespread repressor of IPA (9). Although we found a significant overlap between cisplatin and UV-C regulated IPA events (Fig. S1B), U1 snRNA levels did not seem to be decreased by cisplatin treatment (data not shown).

Our findings also provide new insights into the translation outcome and function of IPA isoforms, especially those produced from the 5’ part of genes (5’IPA isoforms) and more specifically from the annotated 5’UTR part of genes (5’UTR-IPA isoforms). First, while two studies identified 5’IPA isoforms that were degraded in the nucleus, thereby preventing their export and translation (15, 16), our data identify a set of IPA isoforms that are found in the cytosol but are lowly associated with HPs (Fig. 2C) and this set is enriched in 5’IPA, more precisely 5’UTR-IPA (Fig. 2F-H and S2B). Some of these 5’IPA isoforms may have a noncoding function, as shown for the IPA isoform of *ASCC3* (8), which is one of the cisplatin-upregulated 5’IPA (but not 5’UTR-IPA) isoforms that we found to be inefficiently associated with HPs (Fig. 2D-E). Along these lines, a 5’UTR-IPA isoform (called SPUD) with a noncoding function was recently described in the *CDKN1A* gene (14).

Second, while a study identified immune cell-enriched 5’IPA isoforms that have a low coding potential and were therefore classified as lncRNAs (17), their translation status and potential function were not determined, and to date, IPA isoforms were not shown to encode microproteins. Here, we provide evidence that many 5’UTR-IPA transcripts, including those in *PHF20* and *PRKAR1B*, contain sORFs that are translated into microproteins, as indicated by our analyses of Ribo-seq and mass-spectrometry datasets and by our polysome profiling, immunofluorescence and Western blot data (Fig. 4, 6B and S3). Thus, our findings reveal 5’UTR-IPA isoforms as a novel source of microproteins. In the case of *PHF20* sORF#1, the fact that the microprotein is supported by mass spectrometry data but not by our immunofluorescence analysis might be due to diffusion out of permeabilized cells, secretion or instability. Alternatively, the additional sORFs (detected by sORFs.org; Table S11) upstream or 5’-overlapping the sORF#1 of *PHF20* might inhibit its translation in some biological conditions (they might also encode their own microproteins but we have not tagged them).

Third, our data reveal the novel paradigm of miP-5’UTR-IPA genes, that produce -from the same promoter but through an IPA switch- a canonical protein-coding full-length mRNA and a microprotein-coding miP-5’UTR-IPA isoform (Fig. 6A). This paradigm is distinct from the case of sORFs located within mRNAs along with the canonical ORF, as in the case of upstream ORFs (21). Indeed, in miP-5’UTR-IPA genes, the sORF and the canonical ORF lie in distinct transcript isoforms. Such a situation has only been shown for some microproteins encoded by circular RNAs, which are alternative transcripts generated by backsplicing (51).

Our data support the functional relevance of miP-5’UTR-IPA genes. Our data indicate that the miP-5’UTR-IPA isoforms of the *PHF20* and *PRKAR1B* genes are functional, because their depletion decreases cisplatin toxicity (Fig. 3). CRISPR editing of the initiation codon of sORF#2 in the *PRKAR1B* miP-5’UTR-IPA isoform shows that the role of this isoform in cisplatin survival is mediated at least in part by sORF#2 (Fig. 5). The cisplatin-induced relocalization of the microprotein encoded by this sORF (Fig. 4E) also argues for its functionality. In addition, for both the *PHF20* and *PRKAR1B* genes, our data suggest that the miP-5’UTR-IPA and full-length mRNA isoforms have opposite effects on cell growth, and that both miP-5’UTR-IPA isoform expression and full-length mRNA down-regulation may contribute to the anti-proliferative effects of cisplatin (Fig. 3F). Interestingly, the PHF20 canonical protein is involved in the DNA damage response, more specifically in the regulation of TP53 (Tumor Protein P53) protein stability in response to DNA damage (52–54). The potential relevance of the downregulation of the PRKAR1B canonical protein, which is a regulatory subunit of protein kinase A, is less clear, but cyclic AMP signalling has been involved in lung cancer cell sensitivity to DNA damage (55).

Besides miP-5’UTR-IPA, our study identifies other IPA isoforms of potential functional interest. In particular, our finding that cisplatin upregulates a 5’IPA isoform in the coding region of *ZFC3H1* (Zinc Finger C3H1-Type Containing; Fig. S1G), a gene encoding a protein involved in the nuclear degradation of 5’IPA transcripts (15), raises the possibility of a positive feed-back mechanism, whereby *ZFC3H1* transcript truncation in response to cisplatin might enhance the cellular accumulation of 5’IPA isoforms. Likewise, we found that cisplatin upregulates a 5’IPA isoform in the coding region of *INTS6* (Integrator Complex Subunit 6; Fig. S1G), a gene encoding a component of the integrator complex that controls the 3’-end maturation and transcription termination of U snRNAs and the early elongation of specific coding genes; *INTS6* was also involved in the DNA damage response (56–58). Regarding the *PRIM2* IPA isoform identified in this study, it is produced from the coding region of the gene (Fig. 1D) and could be detected in polysomes, but we did not detect a truncated PRIM2 protein by Western blot (data not shown), which might be due to a lack of stability. Nevertheless, our siRNA analyses indicated that this *PRIM2* IPA isoform promotes cell survival to cisplatin (Fig. S5); and the strong decrease of full-length PRIM2 mRNA and protein levels in response to cisplatin (Fig. 1G and S1F) is expected to affect DNA replication, where this protein plays a key role as a subunit of DNA primase. Further studies are required to determine the role of the cisplatin-induced large-scale regulation of IPA:LE isoforms (which is enriched in genes related to the DNA damage response, cell cycle and cell death; Fig. 1C) in cancer cell responses to cisplatin.

In conclusion, our data indicate that upregulation of the IPA:LE isoform ratio is a widespread mechanism of transcriptome regulation by cisplatin, that links different layers of gene expression regulation by cisplatin (*i.e.*, it is impacted by transcription processivity and it impacts the translatome) and is involved in cancer cell responses to the widely used anticancer drug, cisplatin. While little was known about the translation status of IPA isoforms, our findings identify distinct subsets of IPA isoforms that are differentially recruited to polysomes. In addition to 3’UTR-APA that affects 3’UTR length, coding region-APA that typically generates protein isoforms (1–4), and two instances of IPA isoforms with noncoding functions (8, 14), our data also reveal 5’UTR-IPA isoforms that encode microproteins and are functional. The novel paradigm of miP-5’UTR-IPA, that we reveal in this study, implicate in cancer cell response to cisplatin, and find enriched in genes involved in specific functions (*e.g.*, signaling and transcription; Fig. 6C), opens wide avenues of research. Future studies should identify more extensively miP-5’UTR-IPA isoforms (here we focused on cisplatin-regulated ones) and determine their function in physiology and disease.

## DATA AVAILABILITY

The datasets generated in this study have been deposited in the UCSC genome browser (private pre-publication link: https://genome.ucsc.edu/s/clabbe/visuCDDPpaper_Dutertre2021) and in the Gene Expression Omnibus repository (GEO) under accession numbers GSE119978 (token: azazmciqfbatzyd), GSE180854 (token: mnevgiouzfetjoz) and GSE180918 (token: ihmlucmubhyfzil). The complete bioinformatics pipeline for 3’-seq analysis of IPA (3’-SMART package) can be freely downloaded at GitHub (https://github.com/InstitutCurie/3-SMART). Any additional information required to reanalyze the data reported in this paper is available upon request. The plasmids are available upon request.

**Supplementary Data are available online.**

## ACKNOWLEDGEMENTS

We thank Patricia Uguen, Christine Tran Quang, and Albertas Navickas for helpful technical advice.

## AUTHORS’ CONTRIBUTIONS

Experiments were performed by AD, IT, QF, AHM, AC and LT under the supervision of MDu, and by SM under the supervision of BE. Illumina and PacBio sequencing were performed by VR and SL, respectively, under the supervision of SB. Bioinformatics and biostatistics analyses were performed by MC, PG, MDe and CML under the supervision of MDu and NS, and by HM, JBC and NF under the supervision of DA. MDu wrote the paper with help of BE, SV, IT, and AD. All the authors read and approved the final manuscript.

