## Supplemental Figures for "Intronic polyadenylation isoforms in the 5’ part of genes constitute a source of microproteins and are involved in cell response to cisplatin"

### Slide 1
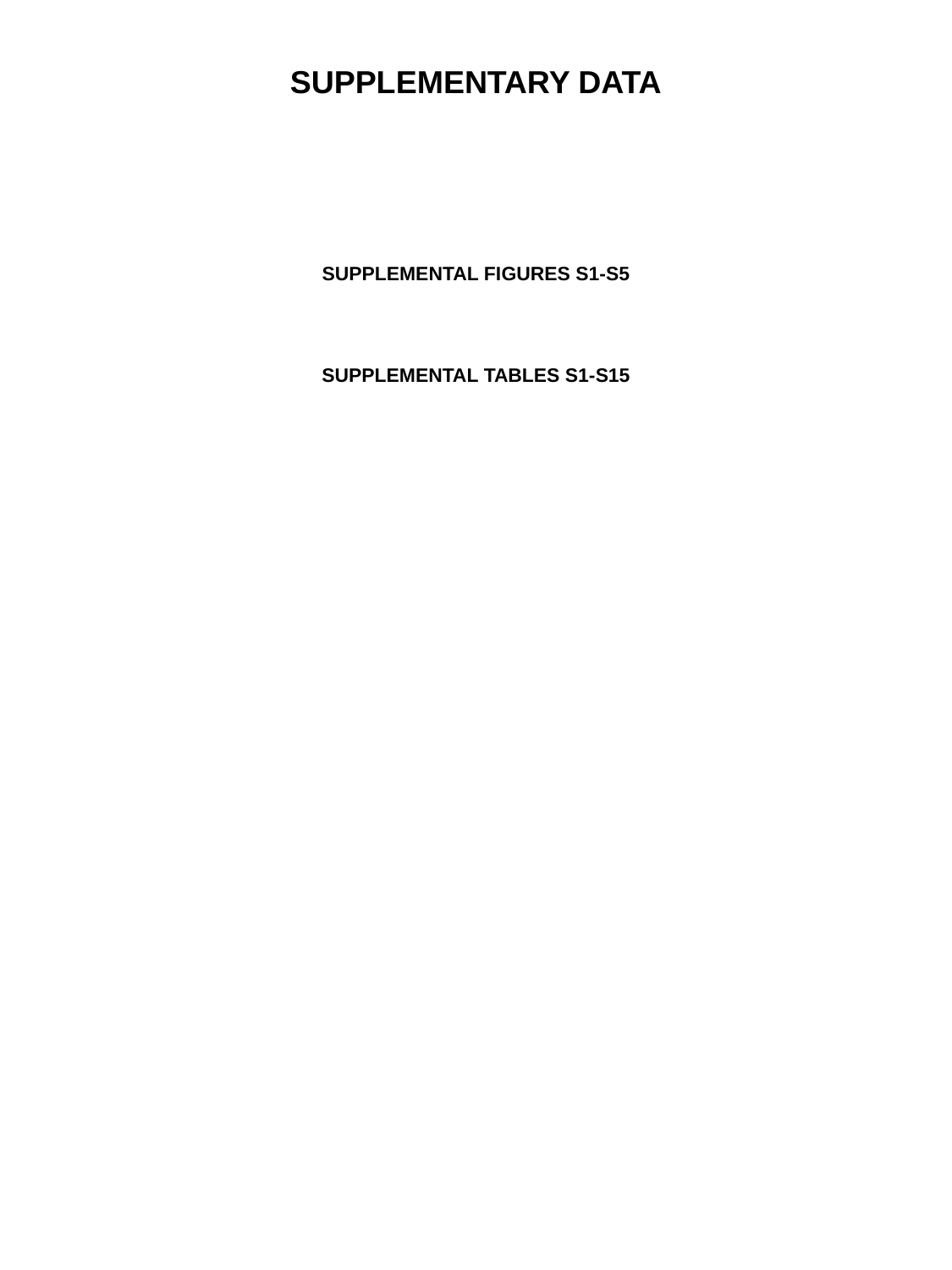

SUPPLEMENTARY DATA
SUPPLEMENTAL FIGURES S1-S5
SUPPLEMENTAL TABLES S1-S15

### Slide 2
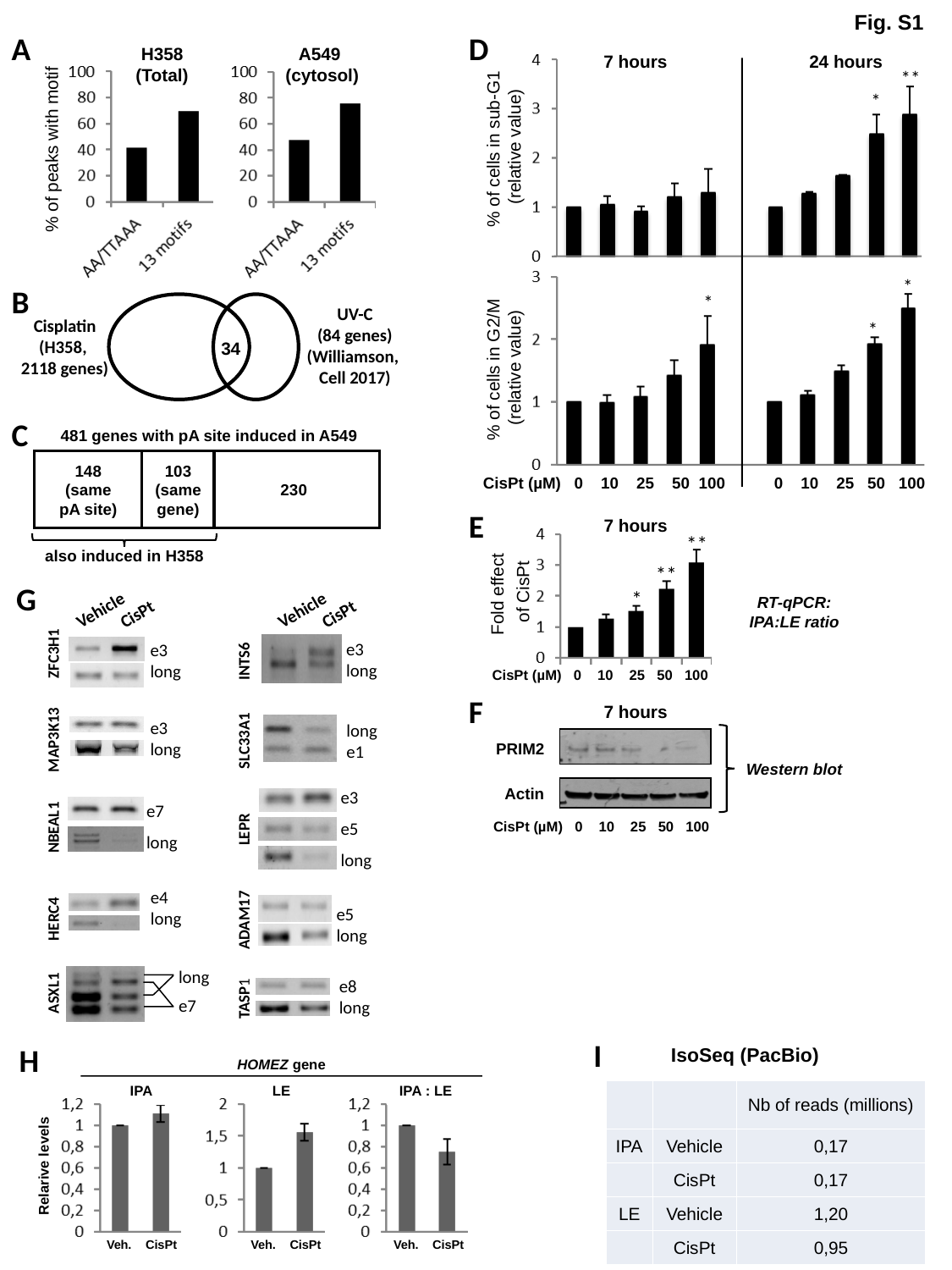

Fig. S1
A
D
H358
(Total)
A549
(cytosol)
% of peaks with motif
7 hours
24 hours
**
*
% of cells in sub-G1
 (relative value)
*
B
*
UV-C
(84 genes)
(Williamson,
Cell 2017)
Cisplatin
(H358,
2118 genes)
*
34
% of cells in G2/M
 (relative value)
C
481 genes with pA site induced in A549
148
(same
pA site)
103
(same
gene)
230
also induced in H358
CisPt (µM) 0 10 25 50 100 0 10 25 50 100
E
7 hours
**
**
Fold effect
of CisPt
Vehicle
CisPt
Vehicle
CisPt
e3
long
e3
long
INTS6
ZFC3H1
e3
long
long
e1
MAP3K13
SLC33A1
e3
e5
long
e7
long
NBEAL1
LEPR
e4
long
e5
long
 HERC4
ADAM17
long
e8
long
 ASXL1
 TASP1
e7
G
*
RT-qPCR:
IPA:LE ratio
CisPt (µM) 0 10 25 50 100
F
7 hours
PRIM2
Western blot
Actin
CisPt (µM) 0 10 25 50 100
I
H
IsoSeq (PacBio)
HOMEZ gene
IPA
LE
IPA : LE
Relarive levels
Veh.
CisPt
Veh.
CisPt
Veh.
CisPt
| | | Nb of reads (millions) |
| --- | --- | --- |
| IPA | Vehicle | 0,17 |
| | CisPt | 0,17 |
| LE | Vehicle | 1,20 |
| | CisPt | 0,95 |

### Slide 3
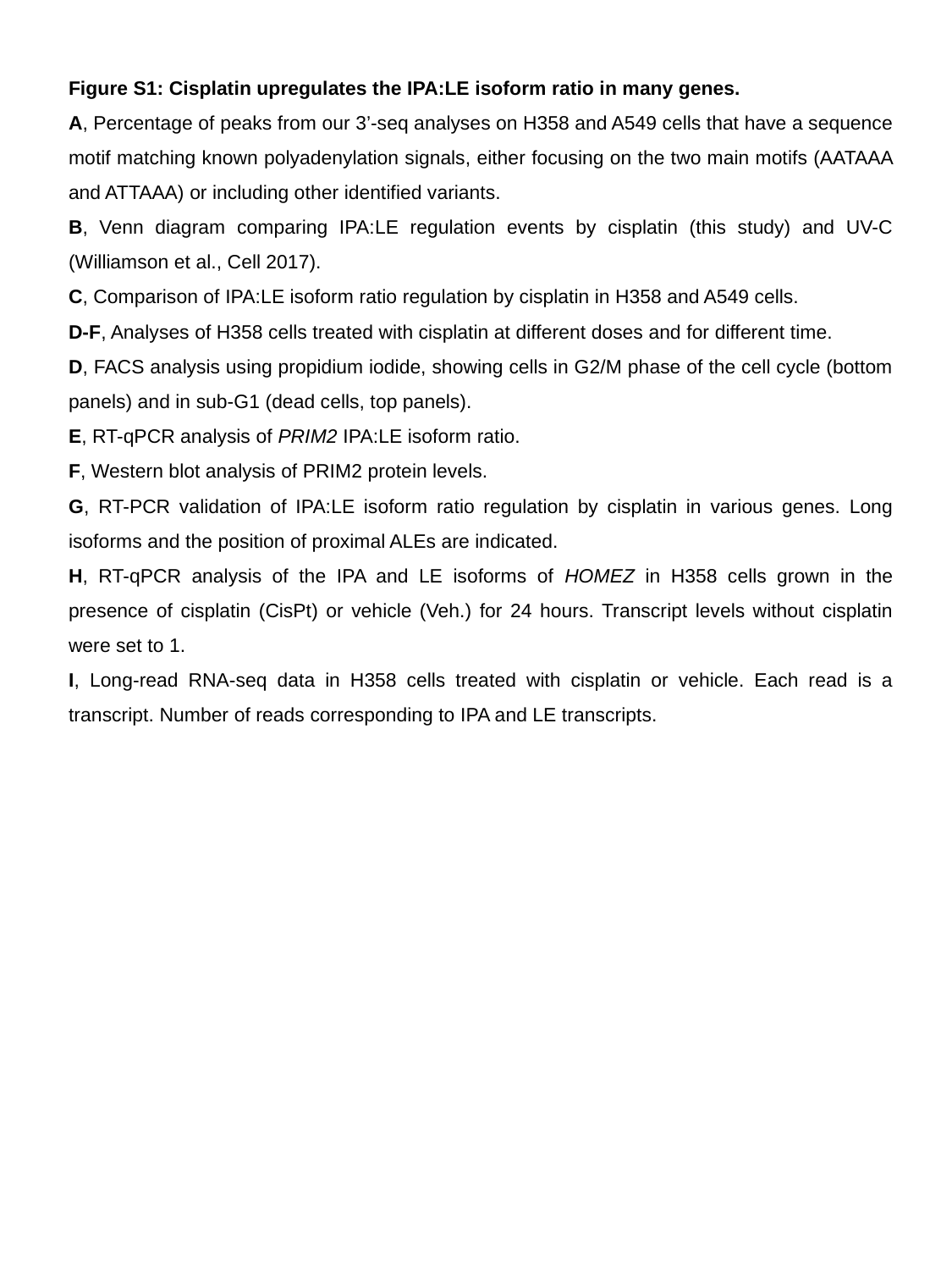

Figure S1: Cisplatin upregulates the IPA:LE isoform ratio in many genes.
A, Percentage of peaks from our 3’-seq analyses on H358 and A549 cells that have a sequence motif matching known polyadenylation signals, either focusing on the two main motifs (AATAAA and ATTAAA) or including other identified variants.
B, Venn diagram comparing IPA:LE regulation events by cisplatin (this study) and UV-C (Williamson et al., Cell 2017).
C, Comparison of IPA:LE isoform ratio regulation by cisplatin in H358 and A549 cells.
D-F, Analyses of H358 cells treated with cisplatin at different doses and for different time.
D, FACS analysis using propidium iodide, showing cells in G2/M phase of the cell cycle (bottom panels) and in sub-G1 (dead cells, top panels).
E, RT-qPCR analysis of PRIM2 IPA:LE isoform ratio.
F, Western blot analysis of PRIM2 protein levels.
G, RT-PCR validation of IPA:LE isoform ratio regulation by cisplatin in various genes. Long isoforms and the position of proximal ALEs are indicated.
H, RT-qPCR analysis of the IPA and LE isoforms of HOMEZ in H358 cells grown in the presence of cisplatin (CisPt) or vehicle (Veh.) for 24 hours. Transcript levels without cisplatin were set to 1.
I, Long-read RNA-seq data in H358 cells treated with cisplatin or vehicle. Each read is a transcript. Number of reads corresponding to IPA and LE transcripts.

### Slide 4
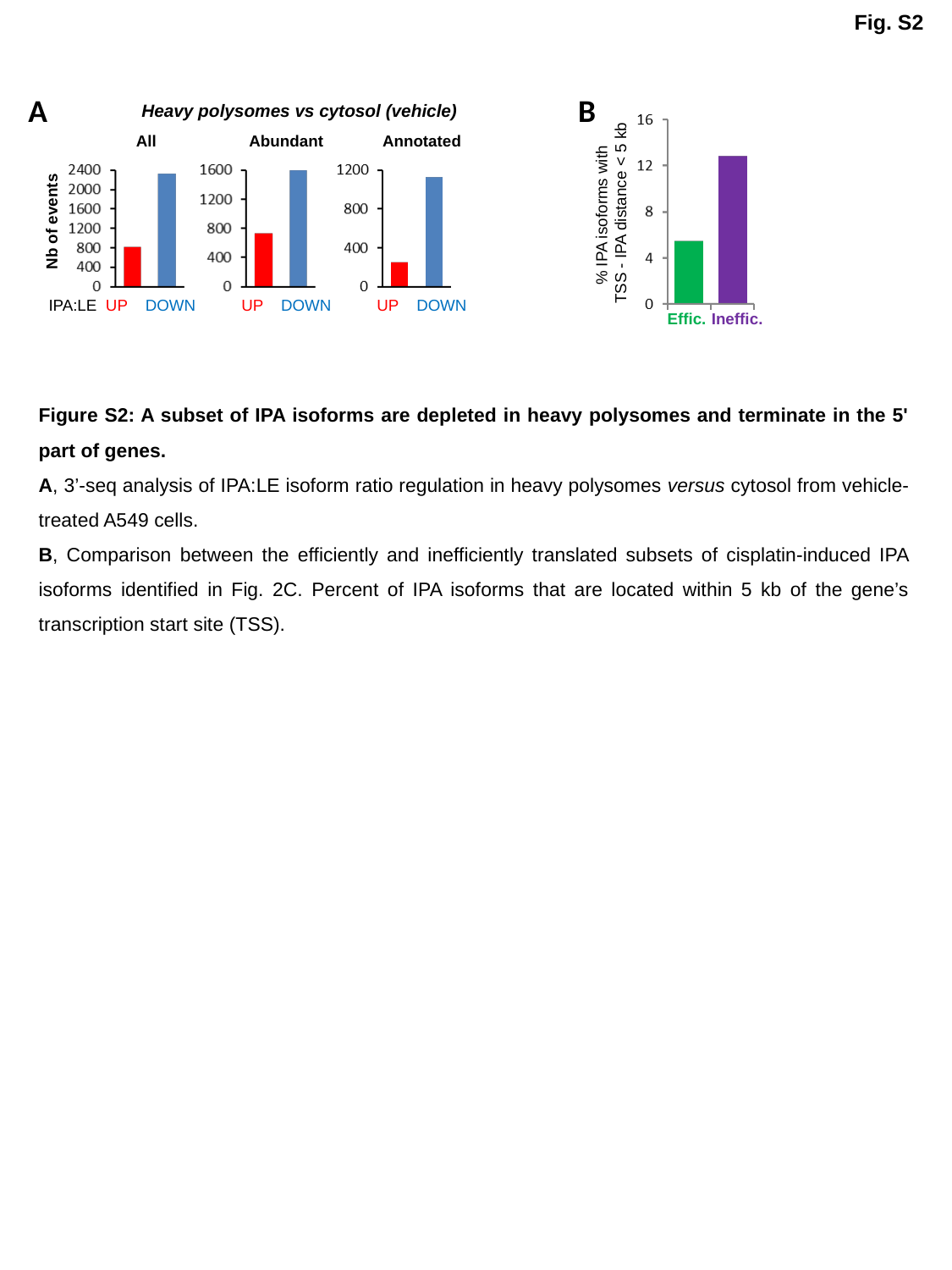

Fig. S2
A
B
Heavy polysomes vs cytosol (vehicle)
All
Abundant
Annotated
% IPA isoforms with
TSS - IPA distance < 5 kb
Nb of events
IPA:LE UP DOWN
UP DOWN
UP DOWN
Effic.
Ineffic.
Figure S2: A subset of IPA isoforms are depleted in heavy polysomes and terminate in the 5' part of genes.
A, 3’-seq analysis of IPA:LE isoform ratio regulation in heavy polysomes versus cytosol from vehicle-treated A549 cells.
B, Comparison between the efficiently and inefficiently translated subsets of cisplatin-induced IPA isoforms identified in Fig. 2C. Percent of IPA isoforms that are located within 5 kb of the gene’s transcription start site (TSS).

### Slide 5
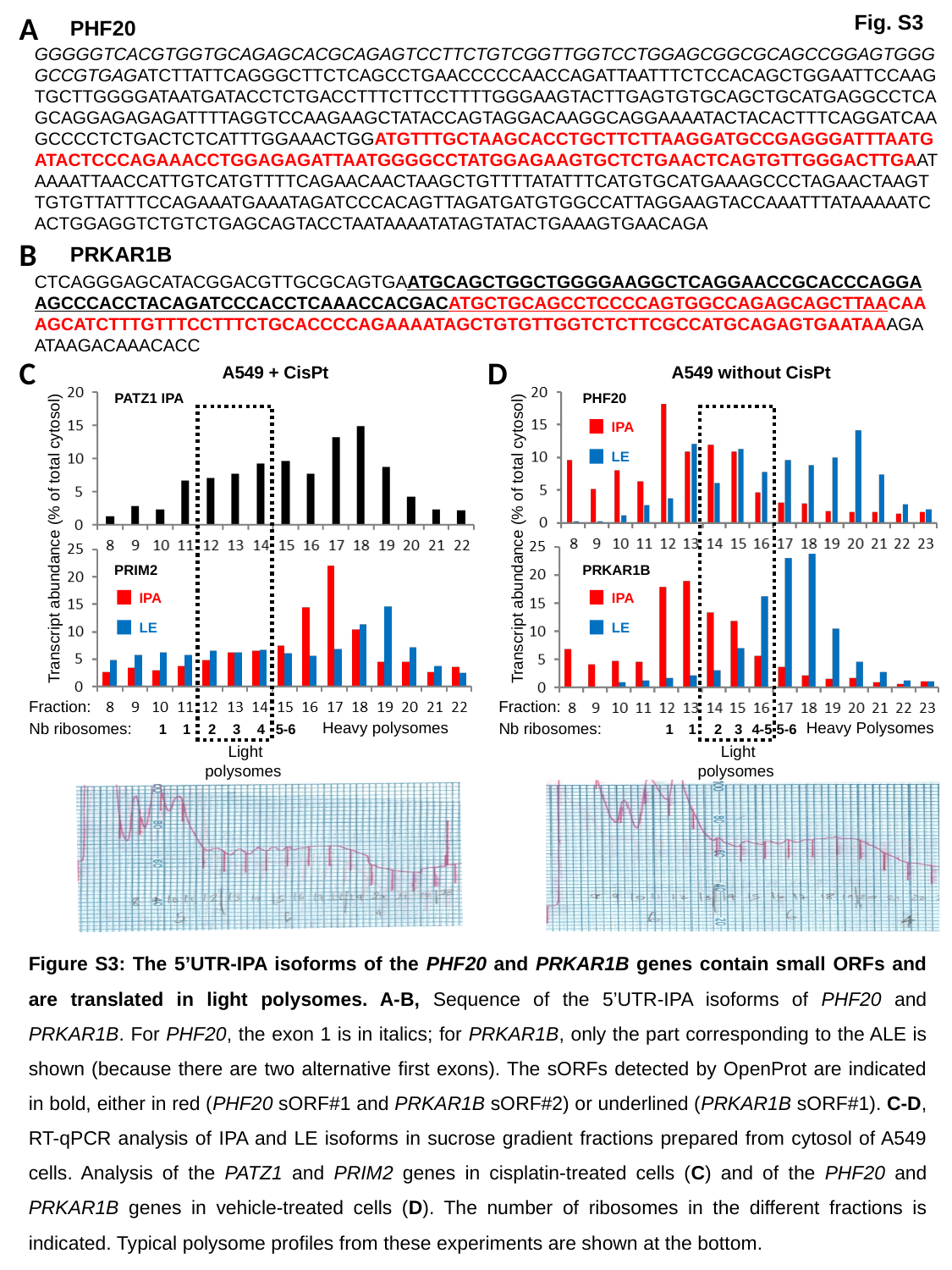

A
Fig. S3
PHF20
GGGGGTCACGTGGTGCAGAGCACGCAGAGTCCTTCTGTCGGTTGGTCCTGGAGCGGCGCAGCCGGAGTGGGGCCGTGAGATCTTATTCAGGGCTTCTCAGCCTGAACCCCCAACCAGATTAATTTCTCCACAGCTGGAATTCCAAGTGCTTGGGGATAATGATACCTCTGACCTTTCTTCCTTTTGGGAAGTACTTGAGTGTGCAGCTGCATGAGGCCTCAGCAGGAGAGAGATTTTAGGTCCAAGAAGCTATACCAGTAGGACAAGGCAGGAAAATACTACACTTTCAGGATCAAGCCCCTCTGACTCTCATTTGGAAACTGGATGTTTGCTAAGCACCTGCTTCTTAAGGATGCCGAGGGATTTAATGATACTCCCAGAAACCTGGAGAGATTAATGGGGCCTATGGAGAAGTGCTCTGAACTCAGTGTTGGGACTTGAATAAAATTAACCATTGTCATGTTTTCAGAACAACTAAGCTGTTTTATATTTCATGTGCATGAAAGCCCTAGAACTAAGTTGTGTTATTTCCAGAAATGAAATAGATCCCACAGTTAGATGATGTGGCCATTAGGAAGTACCAAATTTATAAAAATCACTGGAGGTCTGTCTGAGCAGTACCTAATAAAATATAGTATACTGAAAGTGAACAGA
B
PRKAR1B
CTCAGGGAGCATACGGACGTTGCGCAGTGAATGCAGCTGGCTGGGGAAGGCTCAGGAACCGCACCCAGGAAGCCCACCTACAGATCCCACCTCAAACCACGACATGCTGCAGCCTCCCCAGTGGCCAGAGCAGCTTAACAAAGCATCTTTGTTTCCTTTCTGCACCCCAGAAAATAGCTGTGTTGGTCTCTTCGCCATGCAGAGTGAATAAAGAATAAGACAAACACC
C
D
A549 + CisPt
A549 without CisPt
PATZ1 IPA
PHF20
IPA
LE
Transcript abundance (% of total cytosol)
Transcript abundance (% of total cytosol)
PRIM2
PRKAR1B
IPA
LE
IPA
LE
Fraction:
Fraction:
Heavy polysomes
Heavy Polysomes
Nb ribosomes:
Nb ribosomes:
1
1
2
3
4
5-6
1
1
2
3
4-5
5-6
Light
polysomes
Light
polysomes
Figure S3: The 5’UTR-IPA isoforms of the PHF20 and PRKAR1B genes contain small ORFs and are translated in light polysomes. A-B, Sequence of the 5’UTR-IPA isoforms of PHF20 and PRKAR1B. For PHF20, the exon 1 is in italics; for PRKAR1B, only the part corresponding to the ALE is shown (because there are two alternative first exons). The sORFs detected by OpenProt are indicated in bold, either in red (PHF20 sORF#1 and PRKAR1B sORF#2) or underlined (PRKAR1B sORF#1). C-D, RT-qPCR analysis of IPA and LE isoforms in sucrose gradient fractions prepared from cytosol of A549 cells. Analysis of the PATZ1 and PRIM2 genes in cisplatin-treated cells (C) and of the PHF20 and PRKAR1B genes in vehicle-treated cells (D). The number of ribosomes in the different fractions is indicated. Typical polysome profiles from these experiments are shown at the bottom.

### Slide 6
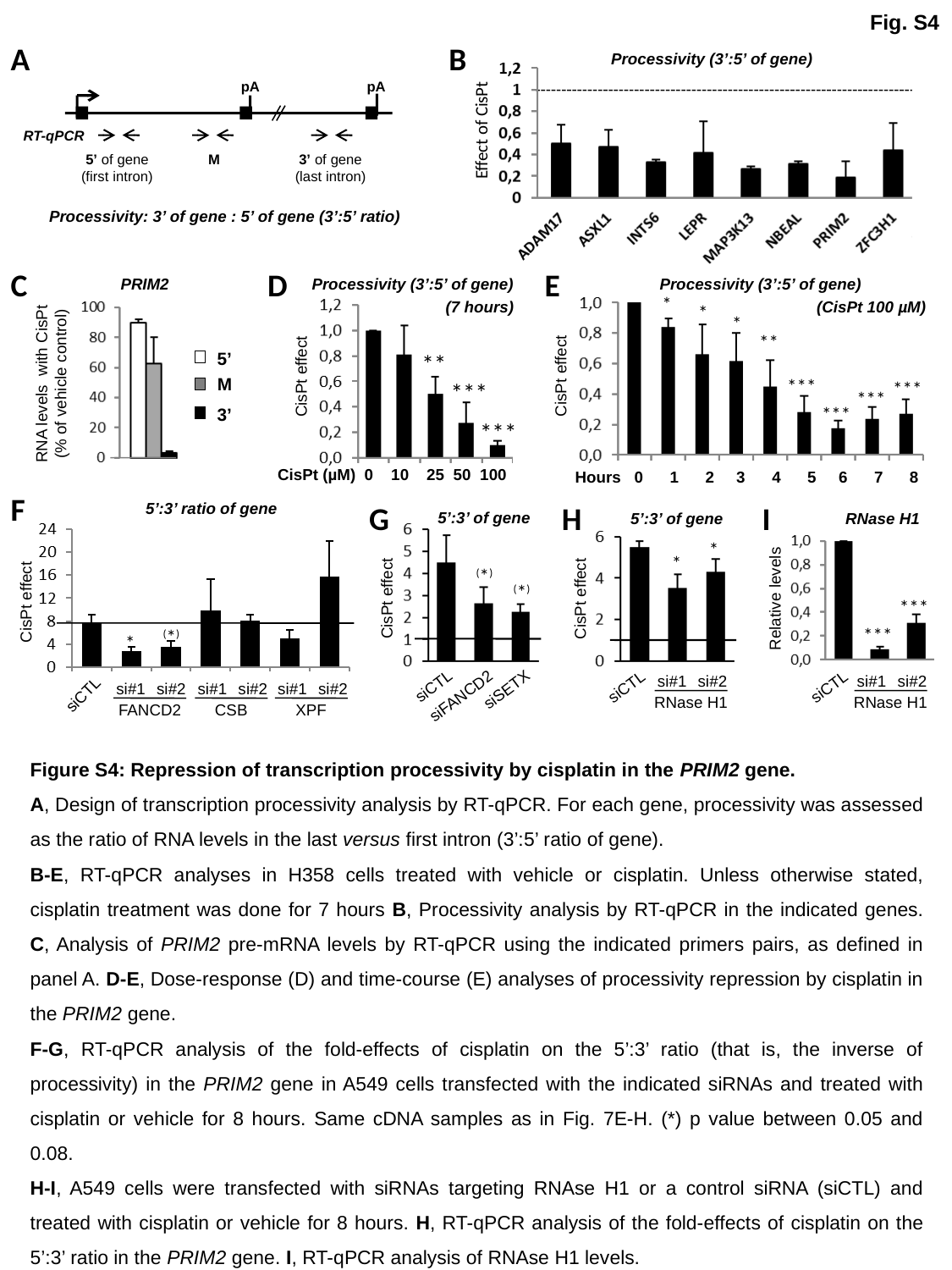

Fig. S4
A
B
Processivity (3’:5’ of gene)
pA
pA
Effect of CisPt
RT-qPCR
5’ of gene
(first intron)
M
3’ of gene
(last intron)
Processivity: 3’ of gene : 5’ of gene (3’:5’ ratio)
D
E
C
PRIM2
Processivity (3’:5’ of gene)
Processivity (3’:5’ of gene)
*
(7 hours)
(CisPt 100 µM)
*
*
**
5’
M
3’
**
RNA levels with CisPt
(% of vehicle control)
CisPt effect
CisPt effect
***
***
***
***
***
***
CisPt (µM) 0 10 25 50 100
Hours 0 1 2 3 4 5 6 7 8
F
G
5’:3’ ratio of gene
H
I
5’:3’ of gene
5’:3’ of gene
RNase H1
*
*
^(^*^)^
^(^*^)^
CisPt effect
Relative levels
CisPt effect
CisPt effect
***
***
^(^*^)^
*
si#1
si#2
si#1
si#2
siCTL
siCTL
siCTL
si#1
si#2
si#1
si#2
si#1
si#2
siSETX
siCTL
siFANCD2
RNase H1
RNase H1
FANCD2
CSB
XPF
Figure S4: Repression of transcription processivity by cisplatin in the PRIM2 gene.
A, Design of transcription processivity analysis by RT-qPCR. For each gene, processivity was assessed as the ratio of RNA levels in the last versus first intron (3’:5’ ratio of gene).
B-E, RT-qPCR analyses in H358 cells treated with vehicle or cisplatin. Unless otherwise stated, cisplatin treatment was done for 7 hours B, Processivity analysis by RT-qPCR in the indicated genes. C, Analysis of PRIM2 pre-mRNA levels by RT-qPCR using the indicated primers pairs, as defined in panel A. D-E, Dose-response (D) and time-course (E) analyses of processivity repression by cisplatin in the PRIM2 gene.
F-G, RT-qPCR analysis of the fold-effects of cisplatin on the 5’:3’ ratio (that is, the inverse of processivity) in the PRIM2 gene in A549 cells transfected with the indicated siRNAs and treated with cisplatin or vehicle for 8 hours. Same cDNA samples as in Fig. 7E-H. (*) p value between 0.05 and 0.08.
H-I, A549 cells were transfected with siRNAs targeting RNAse H1 or a control siRNA (siCTL) and treated with cisplatin or vehicle for 8 hours. H, RT-qPCR analysis of the fold-effects of cisplatin on the 5’:3’ ratio in the PRIM2 gene. I, RT-qPCR analysis of RNAse H1 levels.

### Slide 7
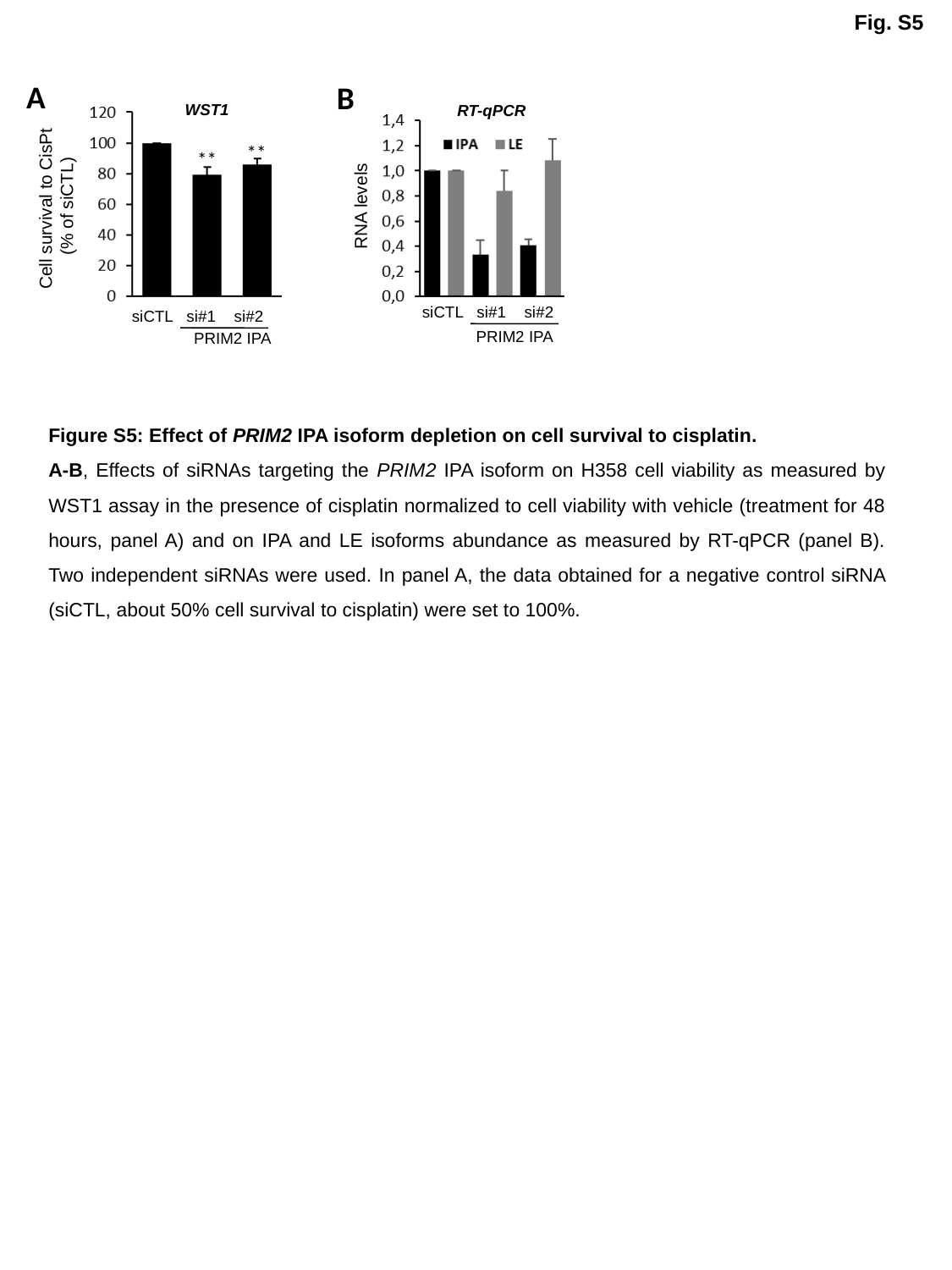

Fig. S5
A
B
WST1
RT-qPCR
**
**
Cell survival to CisPt
(% of siCTL)
RNA levels
siCTL
si#1
si#2
siCTL
si#1
si#2
PRIM2 IPA
PRIM2 IPA
Figure S5: Effect of PRIM2 IPA isoform depletion on cell survival to cisplatin.
A-B, Effects of siRNAs targeting the PRIM2 IPA isoform on H358 cell viability as measured by WST1 assay in the presence of cisplatin normalized to cell viability with vehicle (treatment for 48 hours, panel A) and on IPA and LE isoforms abundance as measured by RT-qPCR (panel B). Two independent siRNAs were used. In panel A, the data obtained for a negative control siRNA (siCTL, about 50% cell survival to cisplatin) were set to 100%.

### Slide 8
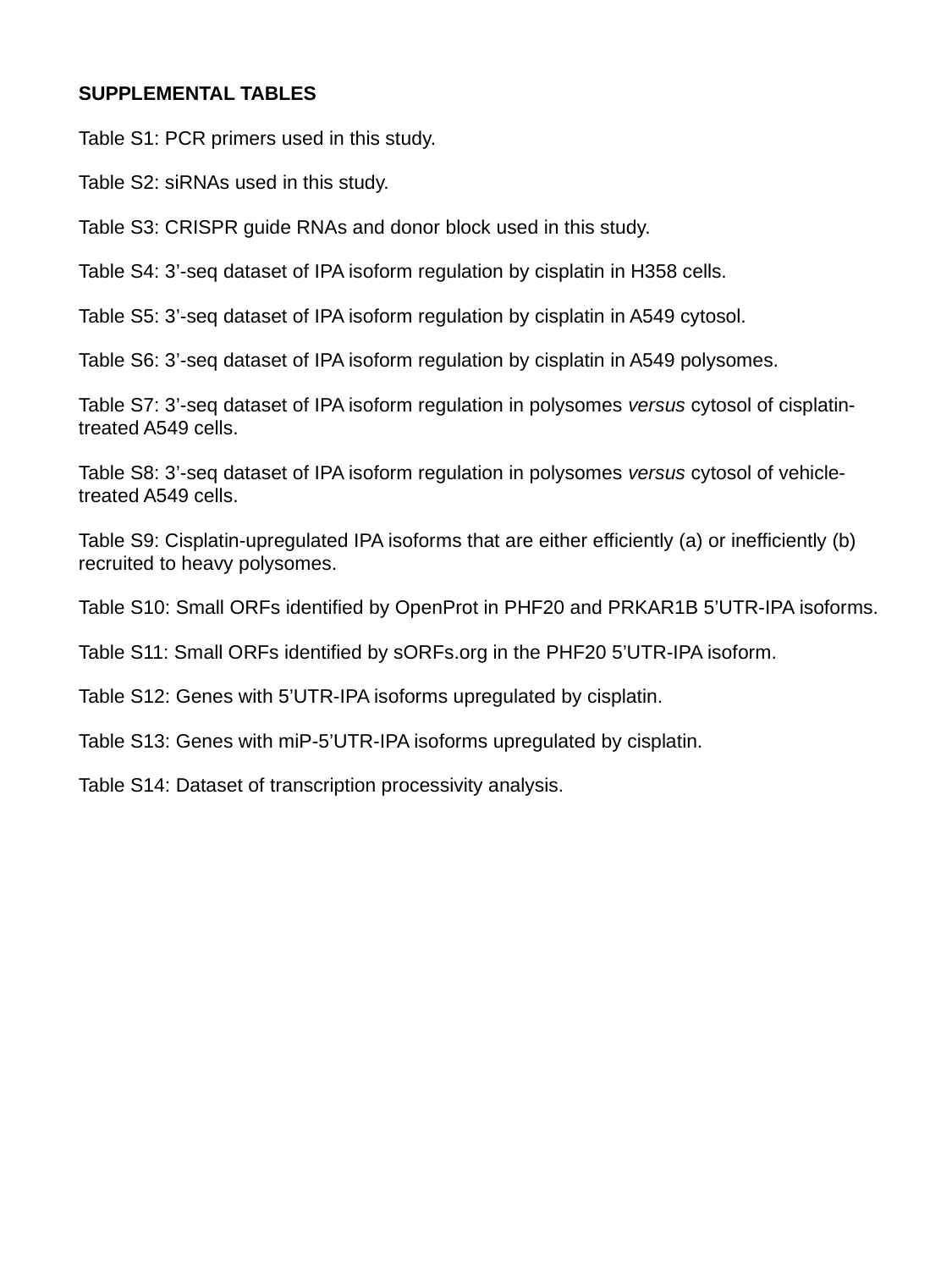

SUPPLEMENTAL TABLES
Table S1: PCR primers used in this study.
Table S2: siRNAs used in this study.
Table S3: CRISPR guide RNAs and donor block used in this study.
Table S4: 3’-seq dataset of IPA isoform regulation by cisplatin in H358 cells.
Table S5: 3’-seq dataset of IPA isoform regulation by cisplatin in A549 cytosol.
Table S6: 3’-seq dataset of IPA isoform regulation by cisplatin in A549 polysomes.
Table S7: 3’-seq dataset of IPA isoform regulation in polysomes versus cytosol of cisplatin-treated A549 cells.
Table S8: 3’-seq dataset of IPA isoform regulation in polysomes versus cytosol of vehicle-treated A549 cells.
Table S9: Cisplatin-upregulated IPA isoforms that are either efficiently (a) or inefficiently (b) recruited to heavy polysomes.
Table S10: Small ORFs identified by OpenProt in PHF20 and PRKAR1B 5’UTR-IPA isoforms.
Table S11: Small ORFs identified by sORFs.org in the PHF20 5’UTR-IPA isoform.
Table S12: Genes with 5’UTR-IPA isoforms upregulated by cisplatin.
Table S13: Genes with miP-5’UTR-IPA isoforms upregulated by cisplatin.
Table S14: Dataset of transcription processivity analysis.
